# Programmed meiotic errors facilitate dichotomous sperm production in the silkworm, *Bombyx mori*

**DOI:** 10.1101/2025.08.03.667510

**Authors:** L Benner, M Richmond, Y Xiang, LS Lee, T Li, Z Yu, D Tsuchiya, S Huang, C Hockens, EC Tromer, RS Hawley, LF Rosin

**Affiliations:** Unit on Chromosome Dynamics, Division of Development Biology, Eunice Kennedy Shriver National Institute of Child Health and Human Development, National Institutes of Health, Bethesda, Maryland, 20982, USA; Stowers Institute for Medical Research, Kansas City, Missouri, 64110, USA; Molecular Genomics Core, Eunice Kennedy Shriver National Institute of Child Health and Human Development, National Institutes of Health, Bethesda, Maryland, 20982 USA; Cell Biochemistry, Groningen Biomolecular Sciences & Biotechnology Institute, University of Groningen, Linneausborg, Nijenborgh 7, 9747AG Groningen, the Netherlands

**Keywords:** Spermatogenesis, meiosis, holocentric, silkworm, eupyrene, apyrene, chromosome segregation, RNA-seq, transcription

## Abstract

The goal of meiosis is typically to produce haploid gametes (eggs or sperm). Failure to do so is catastrophic for fertility and offspring health. However, Lepidopteran (moths and butterflies) males produce two morphs of sperm: nucleated (eupyrene) sperm which fertilize the egg, and anucleated (apyrene) sperm, both of which are essential for fertilization. The meiotic differences in the two types of spermatogenesis have not been well characterized, and our knowledge of the molecular differences between eupyrene and apyrene spermatogenesis are extremely limited in all systems. The only factor identified as being required for apyrene spermatogenesis is *Sex-lethal* (*Sxl*). Here, we show through cytological analysis of early meiotic events that there are several key differences in the genesis of apyrene sperm and eupyrene sperm. Specifically, apyrene spermatocytes fail to condense and pair their chromosomes during meiotic prophase I. In addition, telomeres do not attach to the nuclear envelope. Due to these differences, full-length synaptonemal complex does not form. RNA sequencing of both eupyrene- and apyrene-producing testes revealed distinct changes in transcriptional programs, including down-regulation of a myriad of meiotic genes and cell cycle checkpoint factors during apyrene meiosis. By comparing wild-type and *Sxl*-knockout apyrene testes, we found that *Sxl* is not required for the changes in the expression of the meiotic genes but instead plays a role in checkpoint inactivation to allow this error-prone meiosis to proceed. Together, our findings reveal significant insights into two converging molecular pathways that promote the formation of dimorphic sperm in Lepidoptera.

## Introduction

Dichotomous spermatogenesis, or the production of two distinct yet required spermatozoa, is a phenomenon that occurs in many invertebrate species. This was first described in 1903 through the study of spermatogenesis in both a species of snails and moths (1). More recent work has shown that dichotomous spermatogenesis is a conserved spermatogenic program throughout the order Lepidoptera (moths and butterflies). In the context of Lepidoptera, dichotomous spermatogenesis refers to the production of nucleated sperm (eupyrene) and anucleated sperm (apyrene), with both being required for male fertility (2). The production of these two sperm morphs occurs during distinct timepoints in development. Although slight differences exist in the developmental production of these sperm morphs between species, generally, eupyrene sperm are made first during larval development and the switch to apyrene sperm production occurs during the pupal stages (Reviewed (3)).

How these two different meioses result in maintained or discarded nuclei is unknown, but much of the evidence to date has been acquired through cytological analysis of spermatocytes. Eupyrene spermatocytes have been described as following a conserved, canonical meiotic program (3, 4). In contrast, apyrene spermatocytes lack many fundamental meiotic features. For example, homologous chromosomes fail to pair and the assembly of the synaptonemal complex (SC) does not seem to occur (or is ephemeral in formation), and as a result, bivalents fail to form. Telomeres also fail to associate with the nuclear envelope, further leading to dysregulation of ordered chromosomal orientation that is usually seen during meiosis (Reviewed (3)). The consequence of these events is believed to lead to the disrupted, non-homologous chromosomal alignment during metaphase I. The resulting completion of meiosis I has been characterized by the presence of unresolved chromosome bridges often leading to “fractionation” of chromosomes and the presence of ‘mini-chromosomes’. Daughter cells therefore inherit an unequal chromosome content that becomes further exacerbated in the next meiotic division eventually leading to the total loss of nuclei (Reviewed (3)). These results have garnered the relatively simple hypothesis that the mechanism driving the eupyrene to apyrene spermatogenic switch is the dysregulation of meiotic genes from larval to pupal development (Reviewed (3)).

While the mechanism regulating the eupyrene to apyrene switch is unclear, two distinct factors have been implicated in driving apyrene spermatogenesis. One proposed regulator is hormone signaling, which is known to drive metamorphosis in insects. Evidence for this is that in many Lepidoptera, transplanting larval testes into pupal tissues, regardless of sex, is sufficient to induce the switch from eupyrene to apyrene spermatogenic programs (5). Also, experimentally inducing precocious pupation is sufficient to induce apyrene spermatogenesis earlier than age-matched controls (6). Therefore, there is likely a systemic hormone signal during development promoting the induction of apyrene spermatogenesis. The precise signal is unknown, however, and is commonly referred to as the apyrene secreted inducing factor (ASIF).

The second proposed regulator of apyrene sperm production is the RNA-binding protein Sex-lethal (Sxl). There is a vast amount of literature on the function of Sxl in *D. melanogaster*, where it has been well documented to be the primary regulator of somatic sex-determination, dosage compensation, and female germ cell development (7). In *B. mori*, *S*xl has been shown to be required for apyrene sperm production. Sxl KO (*Sxl-*) testes fail to make functional apyrene sperm and pupal sperm bundles exist in a described ‘intermediary state’, with the failure to eject nuclei that ultimately results in male-specific sterility (8, 9). Despite these findings, the mechanism through which Sxl regulates the apyrene transition, and if there is a relationship with hormone signaling, has not been described.

In our work on eupyrene and apyrene spermatogenesis in the silkworm *B. mori*, we use updated technologies such as Oligopaint DNA FISH and newly described antibodies to *B. mori* meiotic proteins to confirm that apyrene spermatocytes fail to execute a number of conserved meiotic events. Apyrene chromosomes fail to link telomeres to the nuclear envelope, fail to pair homologs, and fail to form a proper SC. To investigate the molecular regulation of apyrene spermatogenesis, we perform RNA-sequencing (RNA-seq) on eupyrene and apyrene spermatocytes and show that the expression of fundamental meiotic genes is globally perturbed during apyrene spermatogenesis, consistent with our cytological data. Finally, gene expression profiling of *Sxl-* testes also provides evidence for a loss of tissue identity in the absence of Sxl, and we find that Sxl is unlikely directly involved in regulating the changes seen in meiotic gene expression between the two sperm morphs. Altogether, our results uncover new cell biology regulating a unique meiotic pathway in moths and provides a testable roadmap for future interrogation of the mechanistic changes that occur between eupyrene and apyrene spermatogenesis.

## Results

### Chromosomes in apyrene spermatocytes fail to decondense, pair, or form a proper synaptonemal complex during prophase I

To investigate telomere dynamics, homolog pairing, and SC formation during eupyrene and apyrene meiosis, we first examined prophase I in larval (eupyrene; Fig 1A, S1A) and pupal (apyrene; Fig 1A, S1A) testes using a combination of immunofluorescence and Oligopaint DNA FISH. As expected for a canonical meiotic prophase I, imaging experiments using a FISH probe labeling the insect telomeric repeat and IF labeling SC proteins revealed that demonstrated that at *B. mori* eupyrene pachytene, telomeres cluster around the nuclear edge (as determined by the edge of DAPI staining) and the SC forms thread-like fibers between homologs throughout the nucleus (Fig 1B top, Fig S1B-C) (4, 10–12). Conversely, in apyrene prophase I, telomeres never cluster and fail to localize to the nuclear edge (Fig 1B bottom). Additionally, consistent with prior reports, we see that a complete SC fails to form during apyrene meiosis (3), with SYCP1 (the transverse filament protein) being essentially absent from nuclei, and lateral element components like SYCP2, SYCP3, and HOP1 being nuclear but not forming classical SC fibers at pachytene (Fig 1B bottom and S2A-B).

**Figure 1.**
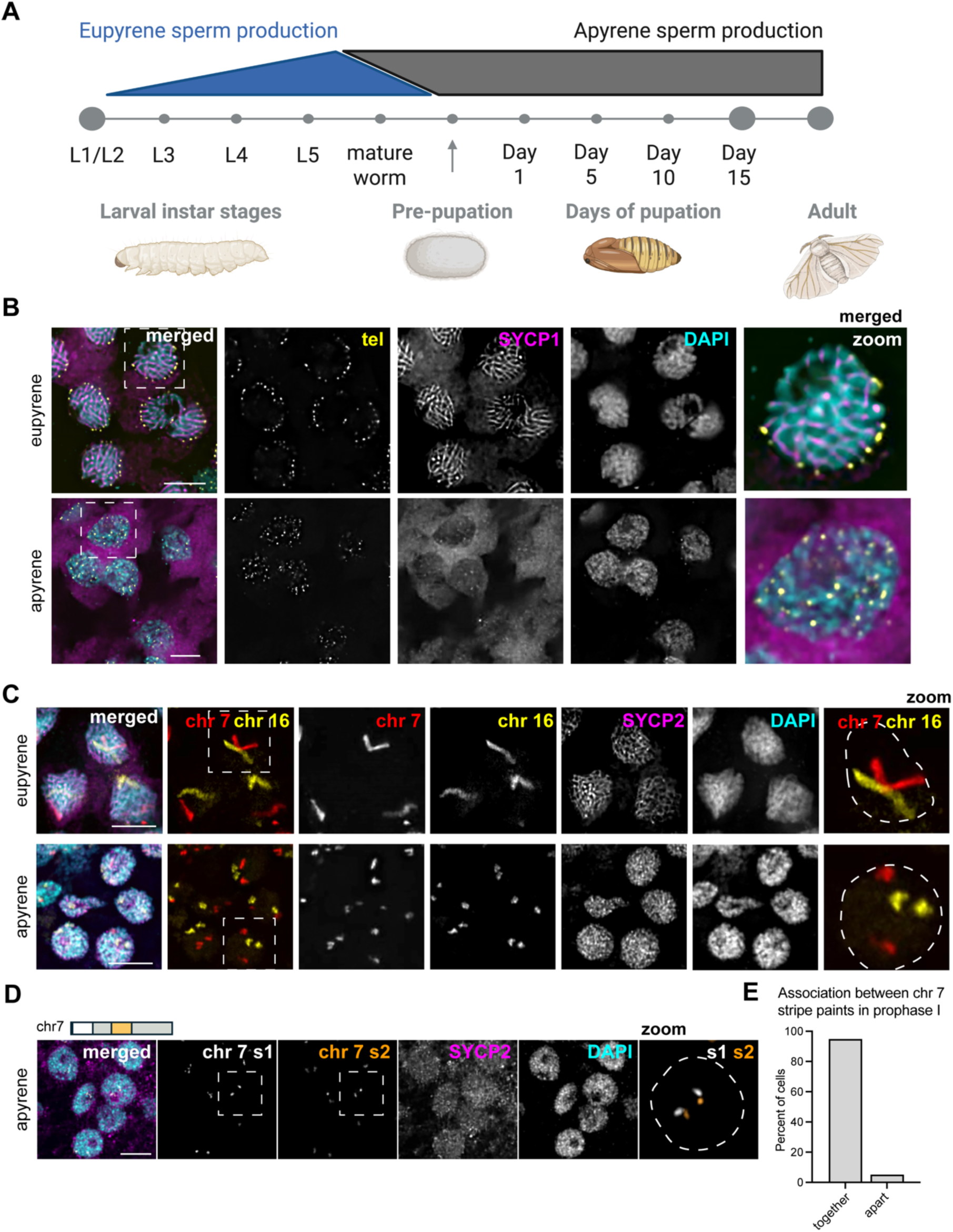
Robust prophase I errors are observed in apyrene spermatocytes. A. Timing of eupyrene and apyrene spermatogenesis during silkworm development. Eupyrene spermatogenesis occurs primarily in larval stages while apyrene development occurs in pupation. A mix of both eupyrene and apyrene production occurs from 5^th^ instar to around pre-pupation (*see Figure S1*). B. Eupyrene (top) and apyrene (bottom) spermatocytes from 5^th^ instar and day 1 pupal testes, respectively. Shown: DAPI (cyan), SYCP1(magenta), FISH to telomeric repeats (tel; yellow). Scale bars = 5 μm. C. Images from cryosectioned slides labeled with Oligopaint FISH for chromosome chr 7 and 16 on prophase I eupyrene (top) and apyrene (bottom) spermatocytes. Unlike eupyrene spermatocytes (top) that form SC and pair homologous chromosomes, apyrene spermatocytes (bottom) do not form proper SC threads and do not pair, as shown through SYCP2 and the Oligopaints. Representative images are shown for spermatocytes spreads using Oligopaints for chr 7 (red) and chr 16 (yellow), co-stained with DAPI (cyan) and SYCP2 (magenta). Apyrene spermatocyte chromosomes are also far more condensed than that of eupyrene spermatocytes in prophase I, and do not display a thread-like shape. Eupyrene images taken from 5^th^ instar larva, while apyrene images are taken from pre-pupa. Scale bar is 5 μm. D. Oligopaint stripes chr 7 s1 (white) and chr 7 s2 (orange) on prophase I cryosectioned apyrene testes (day 1 pupa) demonstrate that each homolog remains intact and does not fragment. This is shown through the striped paints remaining together on each homolog, showing no fragmentation. Co-stained with DAPI (cyan) and SYCP2 (magenta). Image is of apyrene prophase I spermatocytes in day 1 pupa. Scale bar is 5 μm. E. Quantification of chromosome fragmentation as determined by the association of two sub-chromosomal “stripe” paints. Data shown are from day 1 pupae and demonstrate that 95% of all apyrene spermatocytes have their chr 7 s1 and chr 7 s2 associated, showing lack of fragmentation (n = 236).

In agreement with the lack of complete SC, we observed that telomeres never pair during apyrene spermatogenesis. *B. mori* harbor 28 pairs of chromosomes including the sex chromosomes, and thus when telomeres are paired, approximately 56 telomere foci are expected (one for each end). Quantification of the number of telomere foci in prophase I nuclei during apyrene spermatogenesis revealed that more than 80 telomere foci were present in all apyrene prophase I nuclei, while only ∼50 telomere foci were observed in eupyrene prophase I (Fig S2C), supporting the notion that telomeres between homologous chromosomes do not pair in apyrene prophase I. Additionally, chromosome painting with whole chromosome Oligopaints for chromosomes 7 and 16 reveal a complete lack of homolog pairing, as we observed separate FISH signals for each homolog (two FISH signals per nucleus) during apyrene prophase I, in contrast to the single, linear FISH signal observed per nucleus during eupyrene prophase I (Fig 1C and S2D). The irregular, compact shape of chromosomes in apyrene spermatocytes suggests that chromosome decompaction that usually occurs at the beginning of meiosis (4) is absent during this abnormal meiosis. Further support for a lack of chromosome decompaction during apyrene spermatogenesis comes from nuclear size, as we found that eupyrene nuclei are significantly larger than their apyrene counterparts (Fig S2E). While there are clear defects in chromosome organization during apyrene prophase I, chromosomes are not fragmented at the beginning of apyrene spermatogenesis, as confirmed by Oligopaint FISH with sub-chromosomal paints (Fig 1D). Painting of two regions of a single chromosome revealed that chromosomes are almost always intact at the start of apyrene meiosis (Fig 1E). Together, these results indicate a complete lack of homolog pairing at telomeres and along the length of the chromosome during apyrene meiosis I.

### Apyrene metaphase I nuclei harbor unaligned, unlinked univalents

Previous cytological studies on apyrene meiosis in *B.mori* and other Lepidopteran insects have indicated errors with metaphase chromosome alignment and chromosome segregation during anaphase of meiosis I (3, 13–15). To verify these findings, we visualized chromosomes in spermatocytes at metaphase I in both eupyrene and apyrene meiosis. In eupyrene metaphase I, we observed that chromosomes are tightly aligned at the metaphase plate (Fig 2A). Additionally, FISH with chromosome 7 and 16 Oligopaints in eupyrene metaphase I cells showed homologs in bivalent structures as indicated by a single FISH signal per chromosome (Fig 2A). This is in agreement with previous reports in *B. mori* (4) and is consistent with a canonical meiotic program. Conversely, in apyrene metaphase I nuclei, we found severe chromosome alignment defects without a clear “metaphase plate” (Fig 2A), similar to what was observed in prior studies. In line with our analyses of prophase I pairing, FISH with chromosome 7 and 16 Oligopaints in apyrene metaphase I nuclei yielded two distinct FISH signals for each probe, demonstrating that chromosomes form univalent structures at metaphase I instead of being linked in bivalents (Fig 2A-B, Fig S2F). Additional evidence for unlinked homologs at metaphase I comes from chromosome spreads, which showed a median of 26 distinct DAPI bodies at eupyrene metaphase I, but 38 distinct DAPI bodies in apyrene metaphase I (Fig 2C).

**Figure 2.**
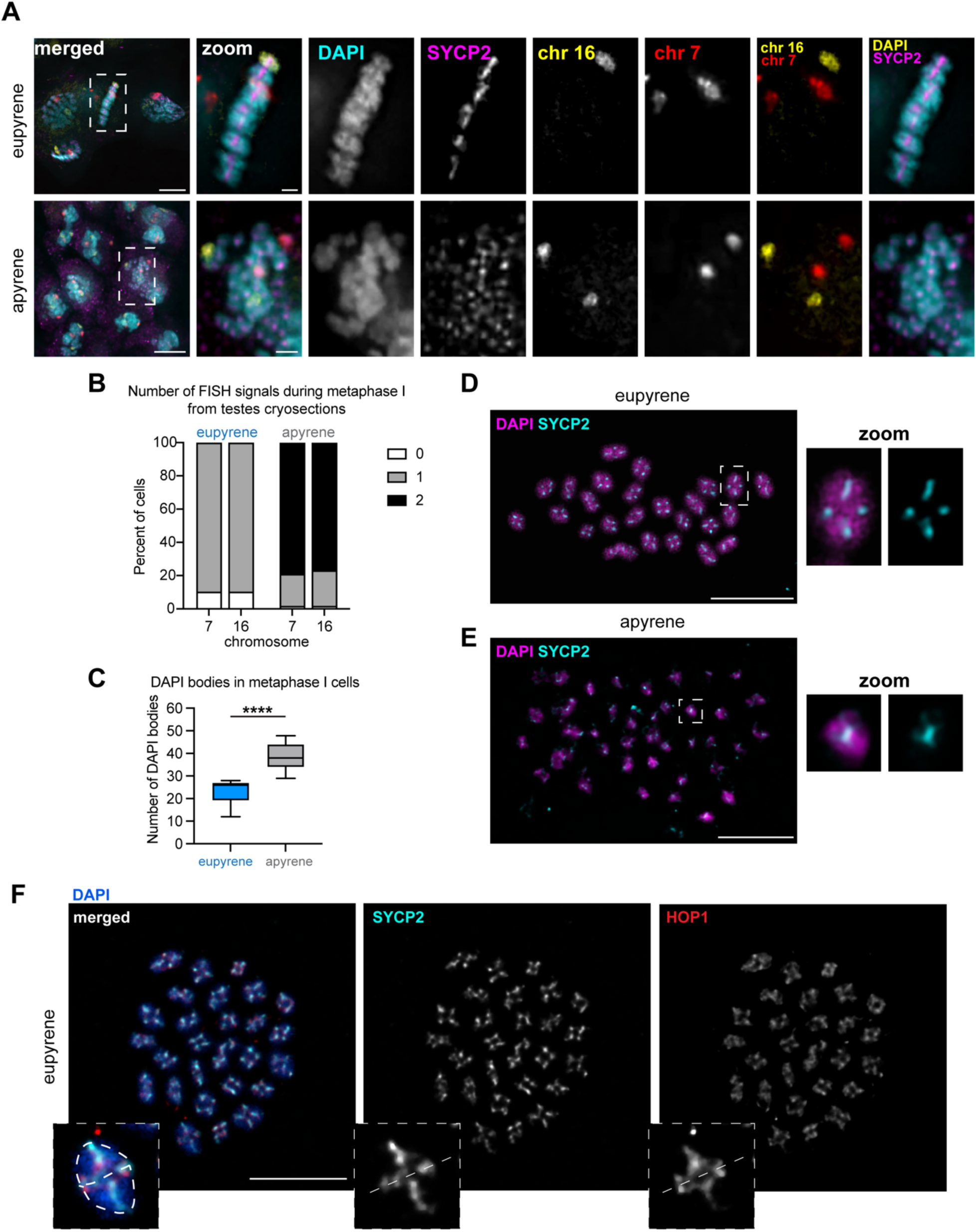
Homologs fail to align and pair at metaphase I. A. Eupyrene (top) and apyrene (bottom) spermatocytes from cryosectioned testes labeled with Oligopaints for chr 7 (red) and chr 16 (yellow), as well as SYCP2 (magenta). Homologs fail to align at the metaphase plate at metaphase I in apyrene nuclei. Additionally, homologs form univalents, and not bivalents, as two distinct FISH signals are observed in apyrene nuclei. Scale bar is 5 μm in merged (left) and 1 μm for zoomed panels (right). Images are maximum-intensity projections of the deconvolved *z*-series though the selected nuclei. B. Quantification of the number of Oligopaint FISH signals in metaphase I cells, as shown in A. 90% of metaphase I eupyrene spermatocytes (5^th^ and mature instars combined) have 1 signal at metaphase I (n = 77). Conversely, 76% & 79% (chr 16 and chr 7) of metaphase I apyrene spermatocytes (day 3 pupa) have 2 signals at metaphase I (n = 103). This indicates that metaphase I pairing follows the same trend as prophase I pairing, with eupyrene spermatocytes successfully pairing while apyrene spermatocytes failing to pair. C. Quantification of the number of chromosome masses in eupyrene and apyrene metaphase I spreads. p<0.0001; were unpaired t-test (Kolmogorov-Smirnov test) between distributions. D. Metaphase I spread from 5^th^ instar larval testes showing that eupyrene spermatocytes have approximately 28 bivalents, and SYCP2 (cyan) remains between homologs as well as between sister chromatids. Right: zoom of indicated chromosome. DAPI is shown in magenta. Scale bar is 10 μm. E. Metaphase I spread from day 7 pupal testes showing that apyrene spermatocytes have more than 28 DAPI bodies, and SYCP2 (cyan) remains between sister chromatids at metaphase I. Right: zoom of indicated chromosome. DAPI is shown in magenta. Scale bar is 10 μm. F. Metaphase I spread from 5^th^ instar larval testes showing that eupyrene spermatocytes have approximately 28 bivalents, and SYCP2 (cyan) and HOP1 (red) remain between homologs as well as between sister chromatids. Right: zoom of indicated chromosome. DAPI is shown in blue. Scale bar is 10 μm.

Likely as a result of chromosome misalignment, we also observed differences in spindle morphology at metaphase I in apyrene meiosis compared to eupyrene meiosis. Apyrene metaphase I is characterized by more bent and narrow spindles, compared to the linear, broad spindles at eupyrene metaphase I (Fig S3). Additionally, the kinetochore protein Dsn1 (part of the Mis12 complex) seems to be broadly mislocalized during apyrene meiosis compared to the distinct kinetochore foci observed on the aligned bivalents in eupyrene meiosis (Fig S3). All of these results are consistent with previous reports of chromosome segregation errors in apyrene meiosis (3, 16), and suggest that this missegregation is likely due to a plethora of errors in early meiotic processes.

### Axial elements of the SC persist through metaphase I in eupyrene and apyrene spermatocytes

Our imaging of eupyrene and apyrene metaphase I spermatocytes not only established that homologs are unlinked at metaphase I in apyrene meiosis, but the images also revealed a curious localization pattern of SC proteins. For example, in our cryosectioned testes, it was apparent that SYCP2 persists on chromosomes at metaphase I in both eupyrene and apyrene meiosis (Fig 2A). To further investigate SC persistence at metaphase, we performed testes spreads followed by immunofluorescence of SYCP1, SYCP2, HOP1, and REC8 (Fig 2D-F, Figs S4, S5). This revealed that lateral components of the SC (REC8, HOP1, and SYCP2) remain associated with chromosomes through metaphase I in both eupyrene and apyrene meiosis (Fig 2D-F, Figs S4, S5). In eupyrene meiosis, where homologs are paired normally, these factors not only localize between homologs at metaphase I, but also between sister chromatids (Fig 2D,F, and Figs S4, S5), explaining their persistence on univalents at apyrene metaphase. Conversely, SYCP1 is gone by metaphase I in eupyrene meiosis (and is entirely absent in apyrene meiosis; Fig S4-S5). This suggests that along with REC8 cohesin, lateral SC elements are involved in sister chromatid associations during both eupyrene and apyrene meiosis I, indicating that SC proteins are being repurposed during *B. mori* spermatogenesis.

### Genes required for homolog pairing and synapsis are downregulated during apyrene spermatogenesis

Our cytological analyses confirmed previous findings that apyrene spermatocytes are indeed failing to properly execute hallmark meiotic events such as telomere clustering, homolog pairing, and metaphase I chromosome alignment. Our data also provide some support for the hypothesis that apyrene spermatogenesis results from a global downregulation of meiotic gene activity, as we observed that SYCP1 protein is not found in the nuclei of apyrene spermatocytes. However, other SC proteins like HOP1, SYCP2, and SYCP3 are present during apyrene meiosis, and they show sister chromatid localization. Still, protein turnover rates might differ between meiotic proteins, and protein presence is not directly indicative of transcriptional changes.

Therefore, we wanted to directly assay meiotic gene expression in eupyrene and apyrene spermatocytes using RNA-seq. To this end, we generated RNA-seq libraries from testes at different developmental timepoints to compare gene expression profiles between the two meioses. We isolated late 4^th^/early 5^th^ instar *B. mori* larvae (>5 cm long) eupyrene testes and late-pupae (∼11-15 days after pupation, after pigmentation had developed in the eyes and abdomen) apyrene testes, as well as the remaining somatic tissue counterparts, and conducted RNA-seq in quadruplicate (Fig 3A-D). As proof of principle, we first checked the expression of genes with previously reported biases in testes or specifically in eupyrene or apyrene meiosis: *vasa* (*vlg*, larval testes-enriched), *wompa* (eupyrene-enriched), and *wimpa* (apyrene-enriched) (17, 18). Consistent with these previous reports, we found that *vasa* has significantly higher expression in larval testes compared to somatic tissues, and *wompa* and *wimpa* are specifically enriched in eupyrene (larval) and apyrene (pupal) testes, respectively (Table S1, S2).

**Figure 3.**
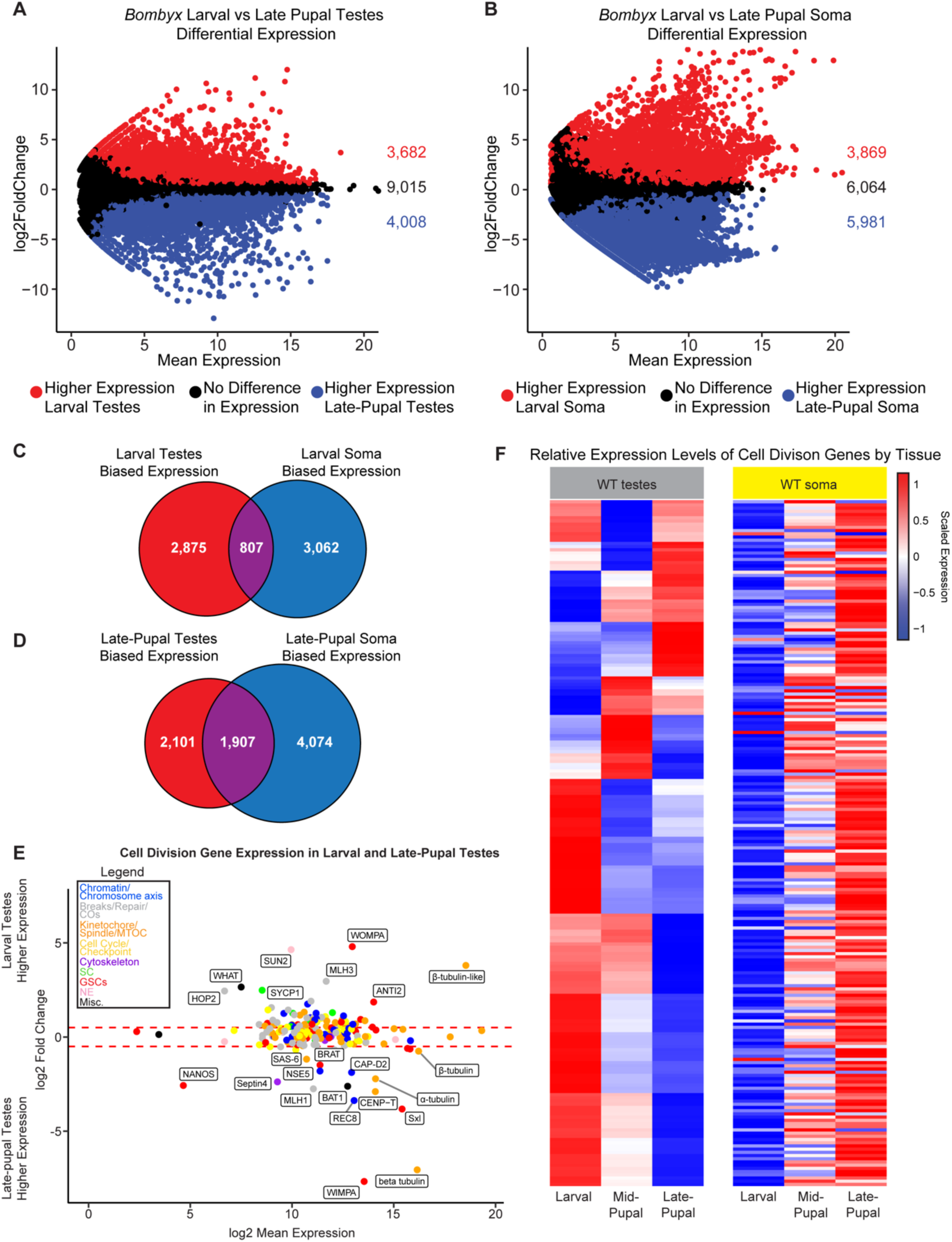
Cell division genes are misregulated during apyrene meiosis. A. MA plot of WT larval vs late-pupal testes gene expression. Red dots indicate genes that are significantly higher expressed in WT larval testes, blue dots indicate genes that are significantly higher expressed in WT late-pupal testes, and black dots indicate genes that are not differentially expressed. B. MA plot of WT larval vs late-pupal somatic gene expression. Red dots indicate genes that are significantly higher expressed in WT larval soma, blue dots indicate genes that are significantly higher expressed in WT late-pupal soma, and black dots indicate genes that are not differentially expressed. C. Intersection of genes that are significantly higher expressed in larval testes vs late-pupal testes and larval soma vs late-pupal soma. D. Intersection of genes that are significantly higher expressed in late-pupal testes vs larval testes and late-pupal soma vs larval soma. E. RNA-seq log2 fold change values of cell division genes between WT larval and late-pupal testes. Select genes and their cell cycle roles, chromatin/chromosome axis (blue), breaks/repair/COs (gray), kinetochore/spindle/MTOC (orange), cell cycle/checkpoint (yellow), cytoskeleton (purple), synaptonemal complex (SC, green), GSCs (germline stem cells, red), nuclear envelope (NE, pink) and miscellaneous (black) are indicated in the legend. F. Heatmap of cell division gene expression (scaled RPKM values) in testes and somatic tissues across development. Genes are hierarchically clustered by their expression in testes tissues.

Within our RNA-seq data sets, we focused on the expression of previously identified *B. mori* meiotic genes (12) as well as newly identified *B. mori* orthologs of meiotic, mitotic, and cell cycle genes (referred to collectively as “cell division genes”) (19, 20) (Table S2). 214 genes in total made up our list of meiotic genes within our RNA-seq analysis (Table S2). Our new gene list includes genes that code for previously unidentified orthologs of critical meiosis proteins in *B. mori*, including the central element SC protein SIX6OS1, several different SUN and KASH domain proteins, and the cohesion-related proteins REC8 and Shugoshin (Table S2). We first looked to see if any of these genes were differentially expressed between larval (eupyrene) and late-pupal (apyrene) testes. We found that 32% (69/214) of cell division genes had significantly higher expression in larval testes than late-pupal testes, while 57% (122/214) of genes were not differentially expressed, and 11% (23/214) of genes had higher expression in late-pupal testes (Fig 3E and Tables S1 & S2). This suggests that there are a number of cell division genes that significantly decrease in expression during the eupyrene to apyrene transition. *Sycp1* had a 5-fold reduction in expression in late-pupal testes (log2 fold change = 2.51, padj = 1.14e^-30^), confirming that the loss of positive antibody staining for SYCP1 in apyrene spermatocytes is correlated with a reduction in gene expression at the late pupal stage. *Sycp2* also showed a greater than two-fold reduction (log2 fold change = 1.28, padj = 1.96e^-19^) and *sycp3* had a greater than 1.5-fold reduction (log2 fold change = 0.90, padj = 2.42e^-14^) in late-pupal testes. While not significantly differentially expressed according to our analysis, *hop1* had a one fold reduction in expression during late-pupal stages (log2 fold change = 0.47, padj = 1.34e^-5^).

Along with SC proteins being downregulated in late-pupal testes, we found that a core component of the LINC complex, SUN domain containing protein 2 (*sun2*) (21), had a striking 25-fold reduction in gene expression in late-pupal testes (log2 fold change = 4.66, padj = 9.07e^-81^) (Table S1). The LINC complex tethers telomeres to the nuclear envelope at meiotic entry and is absolutely essential for proper telomere clustering and the canonical ‘bouquet’ formation of chromosomes during prophase I in many systems (21). Although we do not know with certainty the functional role of SUN2 in *B. mori* meiosis, a reduction in stable LINC complex during apyrene spermatogenesis is highly consistent with the telomere dispersal we observed cytologically. Within our RNA-seq data, we did not find any evidence for dysregulation of other nuclear envelope proteins, such as lamins or nuclear porins. Of the 17 annotated nuclear porin proteins, only 4 showed a slight increase in expression in larval testes while the rest were not differentially expressed. This suggests that nuclear envelopes are likely largely intact during apyrene meiosis, and general nuclear instability cannot explain the cytologically observed meiotic phenotypes.

In addition to determining cell division gene expression changes between larval and late-pupal testes, we also measured cell division gene expression changes between larval testes and ovaries (which are achiasmatic (22, 23)) using RNA-seq. We found that 46% (99/214) of genes were significantly higher expressed in larval testes than ovaries, while 32% (68/214) of genes were not considered differentially expressed, and 22% (47/214) had significantly higher expression in larval ovaries (Fig S6, Table S2). Of the 69 cell division genes that were more highly expressed in larval testes compared to late-pupal testes, 71% (49/69) showed a similar higher expression in larval testes compared to ovaries, indicating that these genes have larval testes-enriched expression when compared to other gonadal tissue types with less canonical meioses. This includes many genes involved in crossover formation, supporting the lack of crossovers in female meiosis.

To rule out the possibility that this downregulation of cell division genes is a consequence of development independent of tissue type, we also looked at the expression of these genes between larval and late-pupal somatic tissues. Strikingly, we found that only 7% (15/214) of cell division genes had lower expression in late-pupal somatic tissues, while 28% (60/214) were not considered differentially expressed. Furthermore, 65% (139/214) had significantly higher expression in late-pupal somatic tissues (Fig 3F). Therefore, we conclude that these cell division genes are not merely globally downregulated during development, and instead, the downregulation of cell division genes in pupae is testes-specific.

### Genes involved in chromosome compaction and axis formation are upregulated during apyrene meiosis

To more broadly investigate changes in gene expression in testes during this developmental transition, we also conducted gene ontology (GO) term enrichment analysis on the set of 3,682 genes that were significantly downregulated in the late-pupal testes compared to larval testes (Table S1). A single biological GO term “meiotic nuclear division”, and two cellular component GO terms, “condensed chromosome” and “synapse” (as in nervous system, not chromosome pairing), were considered enriched (Table S3). This result further supports the idea that there is indeed a global reduction in the expression of meiotic and cell division-related genes in pupal testes.

Although we found that many meiosis-associated genes had a significant decrease in expression in late-pupal testes compared to larval testes, this was not the case for all genes. For example, the meiosis-specific cohesin subunit *rec8* showed a greater than ten-fold increase in expression in late-pupal testes (Fig 3E; Tables S1 and S2). REC8 cohesin forms the basis of chromatin loops during meiosis and is a major component of the SC lateral elements, facilitating chromosome reorganization at the start of meiosis (24). In addition to *rec8*, we also observed increased expression of a condensin I subunit, *capd2*, and the kinetochore protein *cenp-t* in late-pupal testes compared to larval testes (Fig 3E). Condensin I was shown to be required for prophase I cohesin retention in *C. elegans* (25) and CENP-T was previously shown to co-localize with cohesin along the chromosome axis at pachytene in moths (26). Together, the increased loading and stability of REC8 cohesin during prophase I in apyrene meiosis could lead to smaller chromatin loops and hyper-compacted chromosomes, consistent with our observation that chromosomes fail to properly decondense at meiotic entry in apyrene spermatocytes.

Along with axis-associated proteins being increased in pupal testes, we also found a gene annotated as one of two putative *cenp-e* genes (LOC101738413) being significantly upregulated during apyrene meiosis (Tables S1 & S2). Although annotated as *cenp-e*, the associated protein shares only minimal sequence homolog to CENP-E kinesins, and lacks the two conserved CENP-E domains: an N-terminal motor domain and a C-terminal microtubule binding domain (27). A combination of psiBLAST (28) and HMMER (29, 30) produced no definitive orthologs of this gene outside of insects, and HHpred (31, 32) suggests functional domains may include flagella motor function (Table S4). We speculate that this gene may play a role in sperm motility, which is thought to be the essential role apyrene sperm have (3). Importantly, this gene is also not expressed in larval ovary (Fig S6) and thus, we refer to this gene as *bat1* (*Bombyx* Apyrene Testes factor 1; Fig 3E).

### The misregulation of cell division-associated genes in apyrene spermatocytes is largely independent of *Sex-lethal*

The RNA binding protein *Sex-lethal* (*Sxl*) has previously been shown to be required for the eupyrene to apyrene transition in *B. mori* (8, 9). However, the role of *Sxl* in this transition is unknown. To dissect the role of Sxl in *B. mori* spermatogenesis, we first investigated *Sxl* expression in wildtype (WT) tissues. For this analysis, we added a mid-pupal time point (∼5-8 days after pupation) for both the testes and soma to be able to better distinguish between early and late developmental roles of Sxl. Consistent with *Sxl* playing a role in apyrene meiosis, our above RNA-seq analyses show that *Sxl* expression is relatively low in larval testes and significantly increases in expression in both the mid- (log2 fold change = 3.3, padj = 3.9e^-39^) and late-pupal (log2 fold change = 3.8, padj = 1.6e^-82^) testes stages (Fig 3E, 3F, S7). There are two different *Sxl* isoforms found to be enriched in the testes throughout development. The larger *Sxl* isoform has relatively similar levels of protein abundance in both larval and pupal testes. In contrast, the smaller *Sxl* protein isoform showed an increase in abundance towards the end of larval development and remains highly abundant throughout the pupal stages in the testes (8, 9).

Previous reports on *Sxl* expression in *B. mori* suggest that *Sxl* utilizes a single promoter with an alternative splicing decision which results in the large and small protein isoforms (8, 9). However, our RNA-seq data from larval, mid-pupal, and late-pupal testes show evidence for an alternative downstream promoter, which was further confirmed with a *de novo* transcript assembly, that splices into exon 3 of *Sxl* thus encoding a downstream start codon which would be consistent with the smaller Sxl protein isoforms (Fig S7). This promoter is utilized almost exclusively for *Sxl* expression in mid- and late-pupal tissues, consistent with the increase in the smaller Sxl isoform protein abundance during the pupal stages (8, 9).

To determine if *Sxl* is required for the gene expression changes we observed above, we also performed RNA-seq on previously published *Sxl* knockout *(Sxl-) B. mori* late-pupal testes and soma (Fig 4A, C). We first focused on the expression of the above described 214 cell division genes (Table S2), comparing late-pupal *Sxl-* testes to WT late-pupal testes. We found that only 7% (14/214) of cell division genes had higher expression in *Sxl-* testes, while 21% (44/214) had no difference in gene expression, and 73% (156/214) had lower expression in *Sxl-* testes (Fig 4B, Table S2). Among these 73% of cell division genes downregulated in *Sxl-* compared to WT testes were *sycp1*, *sycp2*, and *sycp3*. This finding suggests that *Sxl* is not required for the repression of meiotic genes in apyrene spermatocytes and instead, it might play a role in keeping the minimal levels of meiotic gene expression required for *any* meiotic script to be carried out.

**Figure 4.**
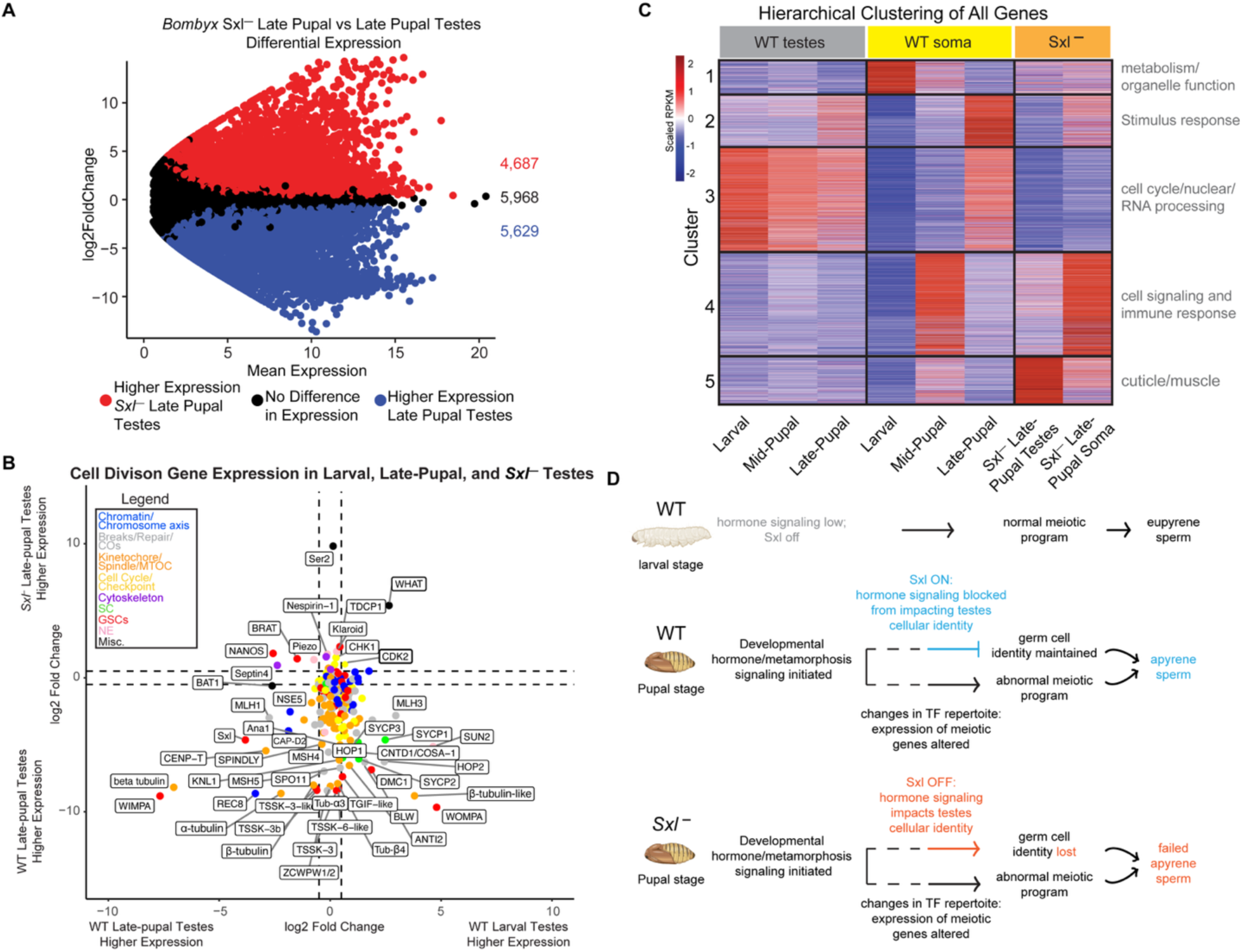
Cell division genes are suppressed independently of Sxl. A. MA plot of *Sxl-* and WT late-pupal testes. Red dots indicate genes that are significantly higher expressed in *Sxl-* late-pupal testes, blue dots indicate genes that are significantly higher expressed in WT late-pupal testes, and black dots indicate genes that are not differentially expressed. B. RNA-seq log2 fold change values of cell division genes between WT larval and late-pupal testes (x-axis) and *Sxl-* and WT late-pupal testes (y-axis). Select genes and their cell cycle roles, chromatin/chromosome axis (blue), breaks/repair/COs (gray), kinetochore/spindle/MTOC (orange), cell cycle/checkpoint (yellow), cytoskeleton (purple), synaptonemal complex (SC, green), GSCs (germline stem cells, red), nuclear envelope (NE, pink) and miscellaneous (black) are indicated in the legend. C. Hierarchical clustering analysis of all expressed genes (scaled RPKM values) for the indicated tissues and developmental timepoints. Five distinct clusters are indicated to the left of the heatmap. Significant representative GO terms for each of the five clusters are indicated to the right of the heatmap. D. Model. Hormone signaling genes are differentially expressed between tissues and time points, and so are global gene expression profiles. We propose a model where differences in hormone signaling lead to differences we observe in putative transcription factors, which regulate either the eupyrene or apyrene spermatogenic pathways independent of *Sex-lethal* (*Sxl*). Expression of *Sxl* is also temporally regulated, and our data suggest that without functional Sxl, the transcriptional profile in testes more closely resembles that of somatic tissue, suggesting a role for Sxl in maintaining germ cell identity. These two pathways cooperatively ensure that the eupyrene-to-apyrene switch occurs successfully.

Interestingly, two genes from our cell division gene list that are significantly increased in expression in *Sxl-* testes compared to WT testes are cell cycle regulators or putative checkpoint proteins. One of these factors, Cyclin-Dependent Kinase 2 (CDK2), is a major cell cycle regulator and had been found to have meiosis-specific roles in prophase I telomere-nuclear envelope tethering and in crossover regulation (33–35). The other gene that is upregulated in *Sxl-* testes compared to WT testes is the second annotated putative ortholog of CENP-E kinesin (Table S2, LOC101739319). However, as with the other putative CENP-E, the associated protein lacks both the N-terminal motor domain and the C-terminal microtubule binding domain. Instead, the protein product of this gene has a central WH2 actin-binding region and several C-terminal Coiled-coil domains but is otherwise predicted to be largely disordered (Fig S8). In some moths, like *Plodia interpunctella*, it also has an annotated Tropomyosin-like domain (Fig S8). The closest ortholog of this protein identified using a combination of psiBLAST and HMMER is *D. melanogaster* SALS (sarcomere length short (36)); Fig S8, and Table S5), which is most highly expressed in muscle cells, but is also lowly expressed in cells from testes and ovaries (37). Functional relationships between SALS and this moth protein are unclear, and therefore, we refer to this gene as *what* (WH2 And Tropomyosin domain). Actin cytoskeletal dynamics play critical roles in meiosis, specifically in prophase I telomere clustering, spindle morphology, and chromosome segregation (38–41). Misregulation of either CDK2 and WHAT during meiosis could lead to meiotic defects, such as the failed telomere clustering and failed pairing phenotypes that we observed in WT apyrene spermatocytes. In agreement with these proteins playing critical roles in meiotic entry, both CDK2 and WHAT are more highly expressed in larval ovary than pupal testes (Fig S9), supporting the idea that they both are required for “normal” moth meiosis. Thus, continual expression of CDK2 and WHAT in *Sxl-* testes could explain why *Sxl-* apyrene spermatocytes cannot fully transition away from a more ‘canonical’ meiotic program.

### Loss of *Sex-lethal* leads to a somatic-like gene expression program in testes

Since we did not have evidence for a global upregulation of cell division genes in *Sxl-* testes, we were interested in looking into other gene expression changes that could help explain why these mutants are unable to make functional apyrene sperm. Globally, we identified 4,687 genes with a significant increase in expression and 5,629 genes with a significant decrease in expression in *Sxl-* late-pupal testes compared to WT late-pupal testes (Table S6). While some of these changes could be due to differences in genetic background of the *B. mori* strains being analyzed, it is also possible that loss of *Sxl* has a dramatic effect on downstream gene expression.

To test this possibility, we performed a K-means hierarchical clustering analysis on all expressed genes in our RNA-seq datasets for different tissues, mutants, and developmental timepoints (Table S6). The heatmap generated from the clustering analysis revealed five distinct gene expression clusters amongst all samples. Interestingly, the global gene expression profile for *Sxl-* testes was not visually similar to any WT testes tissues at any stage, and it seemed that the gene expression profiles were most similar to mid-pupal somatic tissues (Fig 4C). This seemed to also be true for the *Sxl-* late-pupal somatic tissues. To confirm that this was indeed the case, we performed two types of overall similarity metrics that would take into account differentially expressed genes. Principal component analysis, which is a mathematical linear reduction of data into 2-dimensional space, of all RNA-seq replicates showed that *Sxl-* late-pupal testes replicates clustered in closer proximity to mid-pupal somatic replicates and not WT testes replicates (Fig S10). *Sxl-* late-pupal somatic tissue replicates also clustered in closer proximity to WT mid-pupal somatic tissues and not to the developmentally matched WT late-pupal somatic tissues. We also measured the Euclidean distance, a pairwise distance measurement, of all our replicates and found that the tissue with the smallest average Euclidean distance to *Sxl-* late-pupal testes and somatic tissues were also mid-pupal somatic replicates and again, not testes replicates (Fig S10). Overall, this indicates that *Sxl-* testes are more similar in gene expression to mid-pupal somatic tissues than they are to other testes tissues. Also, *Sxl-* late-pupal somatic tissues also show developmentally delayed gene expression profiles. This result suggests that the tissue identity of *Sxl-* testes and somatic tissues is indeed perturbed.

Since we found that *Sxl-* late-pupal testes showed such a wildly different expression profile compared to WT late-pupal testes, we wanted to determine what types of genes were dysregulated in *Sxl-* testes within our clustering analysis. We therefore performed GO term analysis on each gene expression cluster (Fig 4C, Table S6). Cluster 5 genes showed an increase in expression in both *Sxl-* late-pupal testes and mid-pupal somatic tissues. The most significant GO term associated with cluster one genes was ‘structural constituent of cuticle’ which is made up of many of the annotated cuticle proteins and exoskeleton of *B. mori*. In fact, we found that out of the 219 annotated cuticle proteins, 81% (177/219) showed a significant increase in expression in *Sxl-* late-pupal testes compared to WT late-pupal testes.

Another interesting gene expression cluster was cluster 2. Cluster 2 genes showed an increase in expression in WT late-pupal testes and late-pupal soma, but showed a decrease in expression in *Sxl-* late-pupal testes. There were only a few significant GO terms associated with this cluster which were related to stimulus response (Table S7). We also noticed that this cluster showed higher expression in *Sxl-* late-pupal somatic tissues and therefore was the only cluster of genes that showed an appropriate developmental gene expression response. Since this cluster responded appropriately in *Sxl-* somatic tissues, but not testes, this might indicate that this cluster of genes are dependent on *Sxl* activity in the testes but not soma. These results further provide evidence that *Sxl-* late-pupal testes inappropriately express genes that are characteristic of somatic tissues and fail to express genes that are consistent with testes tissues. The identity of *Sxl-* late-pupal testes is clearly perturbed, which may explain their inability to carry out any type of meiotic program.

Within the RNA-seq data, we also identified putative transcription factors (pTFs) in the *B. mori* genome using a recently published deep learning model that is specific for the identification of transcription factors (DeepReg) (42). Out of the 13,437 annotated protein coding genes in the *B. mori* genome, DeepReg identified 2,141 genes as pTFs (Table S8), of which, 2,131 were found to be expressed in the testes according to our RNA-seq analysis. We found that the repertoire of pTFs is significantly different between WT larval and pupal testes, which may contribute to the changes in gene expression we observed between these tissues (Fig S11 and Fig 3). For example, compared to larval testes, 22% (475/2,131) of pTFs are significantly down regulated in late-pupal testes, and 25% (526/2,131) of pTFs become up regulated in pupal testes. In agreement with *Sxl-* testes failing to accurately carry out *any* kind of meiotic program, 68% (1,452/2,131) of pTFs are differentially expressed between *Sxl-* and eupyrene larval testes, and 65% (1,385/2,131) of pTFs are differentially expressed between *Sxl-* and late-pupal apyrene testes (Fig S11).

### Hormone signaling is perturbed in the absence of *Sex-lethal*

Since we didn’t see significant evidence that *Sxl* globally regulates the programmed transcriptional switch between eupyrene and apyrene meiosis, we sought to identify another factor that might be the apyrene spermatogenesis inducing factor (ASIF) (5, 6). Outside of *Sxl*, there is good evidence that hormonal signaling plays a large role in both regulating the onset of meiosis in eupyrene spermatocytes and the switch between the eupyrene and apyrene spermatogenic programs (reviewed in 3). However, the precise hormonal signaling pathway that would promote apyrene spermatogenesis is unclear, and there are likely differences between Lepidopteran species. For example, in *Manduca sexta* and *Cydia pomonella*, ecdysone signaling likely plays the dominant role in regulating meiotic programs (43, 44) while in *B. mori*, findings suggest that a reduction in juvenile hormone signaling during pupation is the resultant trigger (45). Using our RNA-seq data, we wanted to look for changes in hormone-related gene expression that could support the identity of the ASIF in *B. mori*.

To this end, we extracted genes related to juvenile hormone, ecdysone, and other hormone signaling pathways from our RNA-seq data sets and compared their gene expression profiles between development timepoints and tissues (Fig S12). We found different tissue- and time point-specific hormonal signaling transcript profiles; in both testes and somatic tissues, there are only a few hormonal genes that are unchanged across developmental timepoints. Furthermore, most hormonal genes show different developmental expression profiles between the testes and soma. For example, we found that only 22% (11/50) of juvenile hormone genes showed a similar expression pattern in testes and soma at the same developmental time point. For ecdysone signaling related genes, 24% (7/29) showed a similar expression profile across tissues throughout development. This indicates that a vast majority of hormone signaling genes had tissue-biased expression and not developmentally-biased expression. Therefore, at the gene expression level, hormone signaling is highly complex between tissues and development, and there is likely a tissue-specific response to hormone signaling throughout development.

In our *Sxl-* late-pupal testes data, we found that many hormone signaling pathways show an almost opposite gene expression pattern compared to their WT late-pupal testes counterparts (Fig S12). The annotated ecdysone 20-hydroxylase (*Cyp314a1*), which is involved in converting ecdysone into its active 20-hydroxyecdysone (46), as well as the respective ecdysone receptor (*Ecr*), show a dramatic increase in expression in *Sxl-* late pupal testes compared to WT. Similarly, the nuclear hormone receptor E75 and its co-factor hormone receptor 3 (*Hr3*), which have been described to be effectors of ecdysone signaling in *Drosophila melanogaster* (47) and a *Plodia interpunctella* derived cell line (48), are also significantly increased in *Sxl-* testes. However, this pattern of ecdysone-related factors being upregulated in *Sxl-* testes is not universally true. Another described effector of ecdysone signaling, the broad-complex (*Br-c*) (49), is decreased in expression in *Sxl-* testes. Thus, the expression of many hormone signaling genes are perturbed in *Sxl-* testes. Interestingly, we find that overall, the hormonal profiles in *Sxl-* testes are more similar to mid-pupal somatic tissues, consistent with what we found for global gene expression patterns described earlier.

Overall, our data suggest a mechanism where hormone signaling that is initiated during metamorphosis likely triggers multiple changes in the transcriptional programs of testes, including changes in transcription machinery, like pTFs. We propose a model where these changes in pTF profiles alter the expression of testes-specific meiotic genes. At the same time, *Sxl* is turned on, and this blocks hormone signaling from globally impacting testes cellular identity. The combination of these two things leads to testes that produce apyrene sperm. Without *Sxl*, developmental hormones can alter the tissue identity of testes, leading to pupal testes assuming a more somatic-like tissue fate that fail to properly execute the apyrene meiotic program (Fig 4D).

## Discussion

The production of dimorphic sperm in Lepidopteran insects has remained largely an enigma since its discovery in 1902 (1). Here, we identified multiple meiotic pathways that are transcriptionally altered during the formation of apyrene sperm, and these changes in expression correlate with changes in TF abundance in testes. The fact that apyrene sperm morphs are essential for fertility (2, 9) suggests that apyrene meiosis is transcriptionally programmed to be doomed from the start. This model is in agreement with data from the wax moth, *Galleria mellonella*, which also has been shown to have distinct transcriptional programs between apyrene and eupyrene meiosis (18), and the gypsy moth, *Lymantria dispar*, which was shown to have errors as early as apyrene meiotic prophase I (16).

Our cytological and RNA-seq analyses indicate that during apyrene meiotic entry, chromosomes fail to properly form a chromosome axis due to excess REC8-cohesin and Condensin I, and telomeres fail to localize to the nuclear periphery due to misregulation of SUN2 and CDK2. Homologs fail to synapse due to loss of SYCP1 and reduced SYCP2 and SYCP3. The PCH2 checkpoint protein is reduced, allowing for a quick progression through prophase I (50, 51), despite these errors in pairing and synapsis. Finally, chromosomes fail to segregate properly due to all of the prior defects.

The fact that we observed defects at the earliest stages of meiotic prophase in pupal testes could suggest that failed meiotic entry plays a role in apyrene sperm formation. This would mean defects in the differentiation of germline stem cells (GSCs). Supporting this idea, we see that *brat*, which regulates asymmetric cell divisions in flies, including in GSCs (52), is significantly down regulated in apyrene meiosis (Table S1). However, other GSC genes, including many *nanos* orthologs and *pumilio*, are not differentially expressed between our apyrene and eupyrene data sets (Table S1), suggesting that GSCs are at least partially functional in pupal testes.

In addition to characterizing the expression of conserved cell division genes and their roles in eupyrene and apyrene meiosis, we identified *B. mori* orthologs of many conserved meiotic and cell division proteins, as well as two new factors with putative roles in moth meiosis: WHAT and BAT1. Despite both of these factors previously being annotated as putative CENP-E orthologs in *B. mori*, neither contains conserved CENP-E domains, and neither was isolated in previous *B. mori* kinetochore IP-MS experiments (53), suggesting that they are not CENP-E orthologs. We determined that the unannotated gene LOC101743227 is the most likely true CENP-E ortholog. Our computational and transcriptional analyses of WHAT suggest that it is a putative actin binding protein that is reduced in expression during apyrene meiosis. While the exact function of WHAT is still unknown, we speculate that actin-mediated processes could regulate normal chromosome movements or chromosome segregation during meiosis, and thus, reduction of WHAT would lead to defective telomere tethering and spindle/chromosome segregation defects, just as we and others have observed (3, 14, 54). BAT1 is most prominently expressed during apyrene meiosis, and while it has no clear orthologs outside of insects, it has functional domains that suggest a role in flagella motor activity. Since apyrene sperm are critical for sperm motility (3, 9, 55), we suspect that BAT1 could play a major role in this process.

Despite previous findings that *Sxl* is required for apyrene sperm formation (8, 9), our RNA-seq analyses suggest that *Sxl* does not play major roles in downregulating meiotic genes during pupation. Previous imaging of *Sxl-* apyrene sperm bundles showed that the nuclei fail to travel towards the medial region of apyrene sperm bundles and are therefore believed to exist in a transitionary eupyrene-to-apyrene state (8, 9). If an upregulation of meiotic genes is not the cause of failed apyrene meiosis in *Sxl-* testes, then this suggests that there is another pathway dependent on *Sxl* but independent of meiotic gene expression. There is precedent for such a mechanism downstream of *Sxl* in other insects. For example, in *D. melanogaster*, female germ cells mutant for *Sxl* are described to lose their sexual identity and begin to form tumors that show male biased gene expression (56–58). Therefore, *Sxl* is believed to be required for maintaining the sexual identity of female germ cells. Although we are not suggesting that *Sxl* is required for ensuring male-specific identity of testes tissue, our RNA-seq data indicates that *Sxl-* testes are not able to maintain gene expression profiles consistent with WT testes tissue (apyrene or eupyrene). It is possible that one role of *Sxl* is to maintain the germ cell identity of testes tissue and without *Sxl*, pupal testes are now unable to complete apyrene spermatogenesis due to a loss of tissue identity. In support of this model, a recent study found that loss of germ cell identity in *B. mori* testes via mutation in essential germ cell genes *prmt5* or *vasa* leads to a spermatogenic phenotype that highly resembles that of the failed apyrene meiosis in Sxl mutants (17). Loss of germ cell identity via loss of Prmt5 or Vasa function leads to sperm with partial nuclear attrition and partial tail-mediated extrusion (59), exactly as was observed in Sxl mutants (8). The one caveat to this is that it is still currently unknown if *Sxl* is required in the germ cells or somatic cells of the testes. Still, *Sxl* does seem to play a role in allowing apyrene meiosis to occur in WT pupal testes, seemingly by blocking (or bypassing) meiotic cell cycle checkpoints regulated by CDK2 (60, 61) and actin-mediated surveillance (62) via WHAT that allow for the continuation of meiosis in the absence of normal chromosome dynamics.

Our RNA-seq data also suggest that the expression of hormone signaling genes show tissue-specific expression throughout development and each tissue likely has a complex response to systemic hormone signaling carried throughout the hemolymph. The significance of these differences is still unclear. *Sxl-* testes show a perturbed hormonal gene expression response compared to their WT counterparts. Since there is previous evidence that a systemic hormone factor is involved in promoting the eupyrene to apyrene switch (5, 6) and *Sxl-* testes show a perturbed hormonal signaling response, an attractive model for the role of *Sxl* is to integrate the systemic hormonal signaling response specifically within the testes, allowing for the transition from eupyrene to apyrene spermatogenesis. Our RNA-seq data indicate that without functional *Sxl* in pupal testes, there is either a loss of tissue identity or an inability to appropriately respond to hormonal signaling. Either way, the outcome on apyrene spermatogenesis without *Sxl* would likely be the inability to properly execute the apyrene spermatogenic program, consistent with the intermediate phenotype that has been previously described (8, 9).

Curiously, late-pupal somatic tissues show the same upregulation of *Sxl* that is found in the pupal testes. We doubt it is merely a coincidence that the upregulation of *Sxl* in both mid-/late-pupal testes and late-pupal somatic tissues results in similar gene expression profiles. We also found that *Sxl-* late-pupal somatic tissues are also perturbed in their gene expression profiles and are most similar to WT mid-pupal somatic tissues. We are unsure if the gene expression changes in WT late-pupal somatic tissues are a consequence of *Sxl*. However, this result would be consistent with the model that *Sxl* is a mediator of the systemic hormonal signaling that is occurring throughout development. Moving forward, it would be interesting to see if ectopic expression of *Sxl* in somatic tissues earlier in development could alter the gene expression profiles to be more similar to their testes’ counterparts. This would indeed help to further elucidate the role of *Sxl* and its relationship to apyrene spermatogenesis in *B. mori*.

In addition to these findings on the differences between apyrene and eupyrene spermatogenic pathways, our cytological analyses revealed intriguing discoveries regarding protein localization during “canonical” eupyrene *B. mori* meiosis. We found that components of the lateral elements (LEs) are retained at metaphase I in *B. mori* males. In *B. mori* females, Rasmussen described the presence of a large, modified SC that forms between bivalents and persists through metaphase I (22, 23, 63). We recently expanded on this observation and found this structure, now named the bivalent bridge, is an expansion of the LE proteins. As *B. mori* females lack recombination, the bivalent bridge was proposed to act as an alternate method of conjoining bivalents to ensure proper chromosome segregation at metaphase I. The bivalent bridge represents a repurposing of SC components from their canonical roles in synapsis and crossover regulation to directly ensure chromosome segregation. Our finding that LEs are also retained in *B. mori* males intriguingly suggests a repurposing of LE proteins in male meiosis as well. But, since males undergo recombination (64, 65), we wonder what additional role the LE proteins might be playing in this context. Perhaps the retained LEs act as a back-up mechanism to link homologs in the absence of crossovers, as in *B. mori* female meiosis.

Another possibility is that retained LEs play a role in kinetochore function during meiosis. While moths are holocentric during mitosis, we previously showed that during male meiosis, moth kinetochores form discrete, “center-kinetic” foci on bivalents (4, 26). In this meiotic configuration, the kinetochores of sister chromatids are well separated, making it unclear how the cell ensures sister chromatids attach to microtubules from the same pole during meiosis I. In organisms with localized monocentromeres, cohesion is maintained at the centromeres while being lost along the chromosome arms. This cohesion keeps the kinetochores of sister chromatids associated with each other and promotes their attachment to the same spindle pole. Retention of LE proteins, including REC8 cohesin, in *B. mori* males could have been co-opted to help co-orient sister kinetochores.

Importantly, another holocentric organism, *C. elegans*, has been shown to utilize HORMA-domain LEs as part of its mechanism for regulating the placement and retention of the cup-shaped kinetochore and the sequential phosphorylation and loss of REC8 (reviewed in (66)). In this system, differential retention and loss of LEs and other proteins leads to arm-specific, asymmetric loading of pro-kinetochore proteins on the long arms of the bivalent and REC8 being cleaved along the short arm, promoting kinetochore formation on the long bivalent arm. Our observation that LEs remain localized to both long and short arms during *B. mori* male meiosis may indeed suggest a different but similar mechanism, where differences in the length of retained LEs allow the cell to distinguish between the long and short arms of the bivalent to ultimately select which end builds kinetochores. Another possibility is that retention of the LEs provides necessary rigidity to the bivalent so that when one sister kinetochore attaches to a spindle pole, the other sister kinetochore is directed to the same pole. While further experiments will be required to distinguish between these possibilities, the retention of the LE proteins in a pattern distinct from female *B. mori* meiosis is quite intriguing and is a further illustration of the remarkable plasticity in the function of the SC proteins during meiosis.

Something else that remains a mystery is why dimorphic spermatogenesis exists at all. Dimorphic sperm production is not unique to Lepidoptera, and spermatogenesis with both “fertilizing” and “non fertilizing” sperm types has been observed in other insect lineages, such as Diptera, Hemiptera, and Hymenoptera, as well as in gastropods and angiosperms (67). Since these lineages are highly diverged, with non-dimorphic spermatogenesis in intervening lineages, it is likely that dimorphic spermatogenesis evolved multiple, independent times throughout the course of evolution, suggesting that it may confer a selective advantage. Thus, several models have been put forth for why dimorphic sperm production would evolve, including a “nutrition” model, where non-fertilizing sperm provide nutrients to the female or eggs, a “competition” model, where non-fertilizing sperm essentially clog the female reproductive track, blocking competing sperm from reaching the eggs, and a “transport” model, where non-fertilizing sperm assist in transporting the fertilizing sperm through the male and female reproductive tracts. The latter model has the most support (8), but all of these models require further investigation.

In summary, by combining cytological and sequencing-based methods, we find that differences in the dichotomous spermatogenic pathways in *B. mori* initiate early in meiotic prophase I, and our findings reveal significant insights into two converging molecular pathways that promote the formation of the two sperm morphs in Lepidoptera.

## Acknowledgments

We want to deeply thank Cathy Lake and Stacie Hughes for assisting with preparation of and critical reading of this manuscript. We thank Chris Wood for assistance with image analysis, and Ken Campellone for helpful discussion on WH2 domain proteins. We also want to acknowledge Christa Heryanto for help with figures and insect husbandry. Genetic and genomic information was obtained from the UCSC Genome browser. This work utilized the computational resources of the NIH High-Performance Computing Biowulf cluster (http://hpc.nih.gov). *Sxl* mutant silkworm strains used in this study were provided by the National Bio-Resource Project (NBRP) of the MEXT, Japan. R.S.H was an American Cancer Society Research Professor.

## Materials and Methods

### Rearing of silkworms

Wildtype silkworm embryos were obtained from Educational Science https://www.educationalscience.com, or larvae were purchased from FramsChams Chameleon Breeders (https://framschams.com/collections/silkworms). Silkworm embryos were hatched in a petri dish at room temperature (RT). Larvae were fed a diet of fresh mulberry leaves collected in the wild or mulberry chow and feeding boxes were changed regularly. *Sex-Lethal* mutant pupae were obtained from the National Bio-Resource Project (NBRP) of the MEXT, Japan at Kyushu University in Fukuoka, Japan (https://shigen.nig.ac.jp/silkwormbase/about_kaiko.jsp).

### Silkworm Stage Determination

#### For microscopy

There are 5 instar larval stages, with each stage having a molting. Maturing through 1-5^th^ instars takes between 19 to 25 days (68). 5^th^ instar larvae were determined based on their time since hatching and large size. The 5^th^ instar larvae are significantly larger in diameter and length than the previous instars, with maximum pigmentation. Mature silkworm stage proceeds 5^th^ instar stage and occurs right before cocooning. It was determined by a yellowish-translucent color on the ventral side of the silkworm, as well as the silkworms stopping to eat in preparation for cocooning. Pupa stages were determined by cutting a small observation hole in the top of the cocoon. Pre-pupation stage contains a small but normal looking silkworm. After that there is a transition to pupation, where the silkworm will change its appearance to a bulb-like shape that is brown and yellow. Every day after that transition would be marked, hence ‘day 1 pupa’ translates to seeing a pre-pupa the day before, followed by seeing a pupa the next day.

#### For genomics

Larval tissues were harvested from late 4^th^/early 5^th^ instar *B. mori* larvae (>5 cm long), mid-pupae (∼5-8 days after pupation), or late-pupae (∼11-15 days after pupation, after pigmentation had developed in the eyes and abdomen). Eye and abdomen pigmentation was used as the primary marker for late-pupal development to account for potential delays in development in *Sxl* mutants.

### *B. mori* REC8 identification and antibody generation

To identify *B. mori* REC8, the human REC8 (NP_001041670.1) amino acid sequence was used to search the *B. mori* database in NCBI (taxid:7091) using BLAST. This revealed an uncharacterized protein LOC101737592 isoform X1 and X2 [Bombyx mori] (NCBI Reference Sequence: XP_004929454.1) with high sequence homology in the Rad21_REC8 domain to human and Mouse REC8. DNA for the N-terminal 210 aa of XP_004929454.1 was obtained via artificial synthesis by IDT DNA technologies INC in Iowa. The DNA was cloned in pET-21a and the peptide was expressed and purified using the in-frame HIS tag in the vector and Ni-agarose beads (MilliporeSigma). The purified peptide was used to generate antibodies at Cocalico Biologicals in Denver, PA.

### Slide preparation, cell counting, nuclear area size, immunofluorescence, and DNA FISH for Figures 1, 2A-C, S1A and S2

#### Testes processing and cryosectioning

Testes from male *B. mori* at the relevant developmental stage were carefully dissected. For preparation of fixed frozen 5^th^ instar testes samples, the testes were placed in 4% formaldehyde in PBS and incubated at RT for 20 min. Following fixation, the testes were washed in 0.1% PBST (PBS + 0.1% Tween-20) 3 times, 10 min each. The testes were then embedded in OCT compound (Tissue-Tek, Sakura, Japan, cat #4583). For fresh frozen preparation of pupal testes, the samples were promptly embedded in OCT to maintain their structural integrity. The OCT-embedded samples were then snap-frozen on dry ice and stored at −80°C until sectioning.

Cryosectioning was performed using a Cryostat Microtome (Cryostar NX70, ThermoFisher). The cryostat was pre-cooled to −11°C, and the tissue blocks were mounted onto the chuck using OCT with the desired orientation. The blocks were allowed to equilibrate at the cryostat temperature for 30 min to ensure uniform sectioning. Sections of 10 µm thickness were then cut. For fixed frozen testes, sections were collected on Sure Bond Charged Microscope Slides (AVANTIK, cat #SL6332-1) and allowed to air dry for 30 min at RT. For freshly frozen testes, sections were mounted onto pre-chilled Sure Bond Charged Microscope Slides and stored at −80°C for post-fixation before using.

For slides requiring post-fixation, tissue sections on slide were fixed with ice-cold 4% paraformaldehyde (PFA) in PBS for 20 min at RT, followed by three 10 min washes in PBS to remove the fixative. The slides were then transferred to 70% ethanol for 2 hr at −20°C and subsequently incubated overnight in 100% ethanol at −20°C. Afterward, the slides were allowed to air-dry and were then stored at −80°C for future use.

#### Immunofluorescence (IF) on cryosectioned testes

For IF on testes cryosections, an antigen retrieval step was performed prior to immunostaining where the slides were put in 10 mM sodium citrate buffer (pH 6.0) at 90°C for 20 min. The slides were allowed to cool in solution before being immersed into 0.1% PBST for 5 min. Slides were incubated with blocking solution (0.1% PBST + 0.5% BSA) for at least 2 hr before primary antibodies were applied in 0.1% PBST (see Table S9 for concentration) and incubated at 4°C overnight. Slides were washed in 0.1% PBST three times for 5 min before secondary antibody was applied in 0.1% PBST (Table S9) for 2 hr. This was followed by washing in 0.1% PBST three times for 5 min. Slides were stained with 10 µg/mL of DAPI + Vectorshield mounting media for 20 min at RT.

#### IF/FISH for telomeres on cryosectioned testes

To perform IF with fluorescence *in situ* hybridization (FISH), sectioned and spread testes were first treated with 4% PFA for 20 min and then washed three times for 5 min with 0.1% PBST. Prior to immunostaining, an antigen retrieval step was performed where the slides were put in 10 mM sodium citrate buffer (pH 6.0) at 90°C for 20 min. The slides were allowed to cool in solution before being immersed into 0.1% PBST for 5 min. Slides were incubated with blocking solution (0.1% PBST + 0.5% BSA) for at least 2 hr before primary antibodies were applied in 0.1% PBST at 1:250 dilution at 4°C overnight. Slides were washed in 0.1% PBST before secondary antibody was applied in 0.1% PBST (Table S9) overnight at 4°C (12). This was followed by washing in 0.1% PBST three times for 5 min. RNA was digested with 200 µg/mL of RNAse in PBS for 2 hr at 37°C, and slides were rinsed with PBS two times. Slides were dehydrated with ice-cold 100% methanol for 30 min, and the slides were air dried for 1 hr. Telomere probe DNA (100 ng per slide) was mixed with Hybridization buffer (50% formamide, 2X SSC, 1% dextran sulfate, 100 µg/mL salmon sperm DNA), applied on slides, and both genomic DNA and probe DNA were denatured simultaneously on a heating block at 80°C for 5 min. Hybridization was performed at 37°C for 16-20 hr. Slides were washed with 2X SSC for 5 min, 50% formamide/2X SSC for 15 min at 37°C, 2X SSC for 10 min two times, and 2X SSC/0.1% Triton X for 10 min. Slides were stained with 10 µg/mL of DAPI for 30 min at RT. Slides were mounted with ProLong Gold antifade mountant (Invitrogen, cat #36930).

#### IF/FISH with Oligopaints on cryosectioned testes

For IF/FISH with Oligopaints, IF was performed as described above, but with additional wash step following the 5 min wash (PBS + 0.1% triton X-100) after antigen retrieval of 15 min wash with 0.5% PBST (PBS + 0.1% triton X-100). This was followed by FISH protocol as previously described (4, 69). Briefly, cryosectioned slides were post-fixed with 4% PFA for 10 min. After this fix, the slides were washed 2X SSCT for 5 min at RT. This was followed by the slides being treated with 2X SSCT/20% formamide for 10 min at RT, 2X SSCT/50% formamide for 10 min at RT, 2X SSCT/50% formamide for 3 min at 92°C, and 2X SSCT/50% formamide for 20 min at 60°C. Primary Oligopaint probes were resuspended in hybridization buffer (10% dextran sulfate/2X SSCT/50% formamide/4% polyvinylsulfonic acid), placed on slides, covered with a coverslip, and sealed with rubber cement, and let to fully dry. After drying, the slides were then denatured for 3 min at 92°C and incubated overnight at 37°C. The next day, coverslips were removed using a razor blade, and slides were washed as follows: 2X SSCT at 60°C for 15 min, 2X SSCT at RT for 15 min, and 0.2X SSC at RT for 5 min. Fluorescently labeled secondary probes were then added to slides, again resuspended in hybridization buffer, covered with a coverslip, and sealed with rubber cement. Slides were incubated at 37°C for 2 hr in a humidified chamber before repeating the above washes. All slides were stained with DAPI, washed twice for 5 min at RT, and mounted in Prolong Glass Antifade Mount.

#### Staging and identification of prophase I and metaphase I cells in apyrene spermatocytes

For prophase I, staging was determined based on the presence of semi-diffuse SC lateral elements (SYCP2, SYCP3, or HOP1), and nucleus size. Metaphase I cells were identified based on proximity to prophase I cells and the presence of distinct dot-like compacted chromosomes in semi-aligned or cloud-like organizations.

#### Imaging and analysis

Images were acquired with a DeltaVision microscopy system (Leica Microsystems) consisting of a 1X70 inverted microscope with a high-resolution CCD camera (12). Images were deconvolved using SoftWoRx v. 7.2.2 (Leica Microsystems Inc.) software. Image analysis was done using Fiji and cropped in Adobe Photoshop. Brightness and contrast were adjusted minimally and uniformly to visualize signals during figure preparation.

Telomere number was analyzed by the following method: Cell segmentation was performed using Cellpose based on DAPI signal, and a filtering step was implemented to exclude artifacts, debris, and poorly segmented cells. Following segmentation, FISH signal dots within each cell were detected and localized using the Find Maxima function in FIJI, which identifies intensity peaks corresponding to individual FISH signals based on user-defined noise tolerance and prominence thresholds. The number of FISH dots was then counted for each cell to measure the difference.

#### Quantification of spermatocytes throughout development

This was done through the distinctive SYCP2 differences between eupyrene and apyrene spermatocytes, as well as the overall size and shape of nuclei between eupyrene and apyrene spermatocytes using DAPI. SYCP2 appears as distinct foci in apyrene spermatocytes, rather than appearing as threads and therefore making it easily distinguishable from eupyrene spermatocytes. Furthermore, when eupyrene and apyrene spermatocytes are seen side by side, the signal from SYCP2 is much dimmer for apyrene cells, helping to identify them. Similarly, apyrene spermatocytes nuclei are considerably smaller than that of eupyrene spermatocytes, making identification easy using DAPI.

#### Nuclear area determination

Area of the spermatocyte nuclei was calculated using DAPI with all sections of the z-stack used. In ImageJ/Fiji, the background was subtracted using a rolling ball radius of 200.0 pixels with no extra settings checked. Next a Gaussian blur radius of 2.00 was used. The image was then put through the default threshold function to highlight the area of DAPI, and area size overlay was adjusted to the same size as the DAPI signal. Next a size threshold was established using the Analyze Particle function, and a size of 10 µm² to 50 µm² was used for most cell sizes for eupyrene and apyrene. The Measure function was then used to calculate the area for each cell in the image.

### Slide preparation, Immunofluorescence, and DNA FISH for Figure S1B and C

IF/FISH was performed on cytospin-based spermatocyte spreads. Here, testes were dissected out and placed in 0.5% sodium citrate, where they were dissociated by gentle pressure with a pestle. Loose germline cysts were then incubated in the sodium citrate for 10 min. Cysts were spun onto slides using a Cytospin 4 (Thermo) with the following settings: 500 x g, 5 min, max acceleration. Following spinning, cells were fixed to slides with 4% PFA for 10 min, and washed thrice with 0.1% PBS-T. Cells were permeabilized with 0.5% Triton X-100 in PBS for 15 min, blocked in 5% BSA for 1 hr at RT, and then incubated with primary antibodies overnight. The next day, slides were washed thrice in PBS-T, incubated with secondary antibodies, washed again, then post-fixed before proceeding to DNA FISH. The DNA FISH protocol was exactly as previously described (70).

Slides were imaged on a Leica DMi6000 inverted wide-field fluorescence microscope equipped with an APO 63x/1.40 Oil objective (Leica), Leica DFC9000 sCMOS Monochrome Camera, EL6000 light source, DAPI/FITC/CY3/CY5 filter cubes, and LasX software. Images were deconvolved using Huygens deconvolution software (SVI) and tiffs were generated with imageJ.

### Slide preparation and Immunofluorescence Figures 2D-F, S3, S4, and S5

For IF on manual spermatocyte spreads, testes were harvested from 5^th^ instar *B. mori* larvae (>5 cm long) or mid-stage pupae (∼6-10 days after pupation) in 1X PBS, washed twice in hypotonic buffer (0.1 M sucrose, 5 mM EDTA, 0.1 mM phenylmethylsulfonyl fluoride (PMSF)), and incubated in fresh hypotonic buffer for 30 min on ice. Testes were then homogenized in the hypotonic buffer with a pestle and the spermatocyte suspension incubated for 30 min on ice. Spreads were fixed (1.6% PFA, 0.15% Triton-X, 0.05 M sucrose, 5 mM EDTA) in a humidified chamber for 3 hr, and then air-dried at RT. Slides were then washed in Photoflo for 2 min before proceeding to IF. For IF of SC components, immediately following washing, slides were blocked in 5% milk in 0.1% PBST for 1 hr at RT. For DSN1 and Tubulin staining, cells were first subjected to permeabilizations with 100% methanol at RT for 20 min, then PBS-Triton 0.5% at RT for 15 min before blocking in 2% BSA at RT for 1 hr. For all slides, primary antibodies were diluted in blocking solution to the concentrations listed in Table S9 and incubated on slides under parafilm coverslips overnight at 4°C. Slides were then washed thrice in 0.1% PBST before adding secondary antibodies for a 1 to 2 hr RT incubation. After washing off secondary antibodies, slides were DAPI stained, washed thrice in 0.1% PBST, mounted in Slow-Fade or Prolong Diamond, and sealed with nail polish before imaging.

Images in Figures S4 and S5 were acquired on a Leica DMi8 widefield inverted fluorescence microscope equipped with APO 63x/1.40 CS2 oil objective (Leica), K8 CMOS Monochrome Camera, LED8 light source, and with DAPI/FITC/CY3/CY5/CY7 filter cubes. All images were processed using the LasX software with Leica Thunder Deconvolution and tiffs were created in ImageJ. Confocal images in Figure S3 were acquired on a Leica Stellaris 8 Resonant Scanning Confocal with 4 HyD S detectors and 1 HyD X SP detector, and SuperZ Galvo stage. Confocal images were post-processed with Huygens X11 Deconvolution software (SVI).

### RNA-seq: Tissue collection, library generation, and analysis

Gonads and somatic tissues (whole carcass minus gonads) for RNA-seq were harvested from WT *B. mori* males and females, staged as described above. *Sxl-* late-pupal testes and somatic tissues were also collected (∼11-15 days after pupation, after pigmentation had developed in the eyes and abdomen). All tissues were homogenized in TRIzol and total RNA was isolated using the Direct-zol RNA Miniprep Plus Kit (Zymo Research, Tustin, CA) following manufacturer’s instructions. rRNA depleted RNA libraries were generated using Zymo-Seq RiboFree Total RNA Library Kit (Zymo Research) and indexed with Zymo-Seq Unique Dual Index (UDI) Primer Plate (Zymo Research) according to the manufacturer’s instructions. Libraries were sequenced on a NovaSeq 6000 SP200 flow cell (100 bp paired-end sequencing, Illumina, San Diego, CA). Biological replicates were done in quadruplicates except for one larval ovary replicate because it was a failed library and therefore removed from the analysis. RNA-seq reads were mapped to the NCBI *B. mori* genome (NCBI RefSeq assembly:GCF_030269925.1) (71) with HISAT2 software (72) (-k 1 --rna-strandness RF --dta). Mapped reads were then sorted with samtools (73) and then HTseq-count (74) (-f bam -i gene_id -s reverse -r pos) was used to count the number of reads mapping to each gene utilizing the NCBI *B. mori* GTF annotation file (NCBI RefSeq GTF:GCF_030269925.1). DeSeq2 (75) was used to determine differentially expressed genes between developmental timepoints, tissues, or genotypes. A log2 fold change value greater than 0.5 or less than -0.5 with a p-adjusted value less than 0.05 were the cutoff parameters for determining if genes were differentially expressed or not between samples. For *de novo* transcript assembly, stringtie software (76) was used to create individual GTF files from each RNA-seq replicate’s bam file using the NCBI *B. mori* GTF file as a reference annotation. Individual GTF files were then merged with the stringtie merge function to create a single *de novo* transcript assembly. BigWig tracks were generated with Deeptools software bamCoverage (77) (-bs 5 --effectiveGenomeSize 530000000 --normalizeUsing BPM). Gene level tracks were visualized with UCSC Genome Browser (78). ggplot2 was used to generate MA plots (79) and pheatmap was used to generate heatmaps (80). RNA-seq replicate PCA analysis was performed with R software (81) plotPCA() and euclidean distances for hierarchical clustering analysis was performed with the dist() function. Gene level hierarchical clustering analysis for all tissues was performed with the kmeans() function (k=5). Gene ontology analysis on differentially expressed genes and clusters was performed with g:Profiler’s g:GOSt functional profiling software (82). The *B. mori* (*B. mori* (Domestic silkworm, p50T)) organism was used as the background gene set and a p value threshold of 0.05 was used to determine functional gene ontology terms.

### Protein homology searches and cell division gene validation

For annotation of genes potentially involved in cell division (Table S2), we collated a list containing: 1. cell cycle regulators (APC/C, cyclins, CDKs), 2. general germ cell and germline stem cell factors with a focus on those found in *D. melanogaster* and *B. mori* (e.g. piRNA pathway genes, spermatogenesis, stem cell markers), 3. kinetochore proteins (including spindle assembly checkpoint (SAC)), 4. chromatin architecture proteins (e.g. insulators, cohesin, condensin and SMC5/6), 5. known conserved meiotic proteins (83), and 6. an updated version of the genes from the “meiotic toolkit” (12). Genes in groups 5 and 6 defined above include genes involved in DNA double strand break formation and repair (DSBs), homologous recombination (HR), synaptonemal complex formation (SC), pro-crossover factors (CO), and a number of meiotic kinases. To search for orthologs, we employed a multi-layered strategy. We first selected relevant Hidden Markov Models (HMMs) of orthologs of our gene set from the EggNOG (KOGs/ENOGs – EggNOG mapper v2) (84) and Panther (85) database using the automated annotations in Uniprot (86) for both *H. sapiens* and *D. melanogaster*. In addition, we collected HMMs from previous efforts to detect specific gene families for kinetochores (87), chromosomal architecture (88), and the meiotic toolkits mentioned above.

Candidates were subsequently recovered using HMMsearch from the HMMER package with standard settings (89). Significant hits were cross-validated for orthology by performing reciprocal similarity searches between *B. mori* and *H. sapiens*/*D. melanogaster* via direct BLASTp using the bidirectional-best-hit principle (90). For large gene families with many paralogs (e.g. kinases), we generated phylogenetic trees to delineate orthologs using the IQ-tree webserver (standard settings including model selection and 1000 UF bootstraps) (91).

In case no homology links could be established through our standard approach, we used three remote homology detection routes: (1) HMMsearch without heuristic filtering (‘–max’ setting) that occasionally finds remote orthologs at the cost of high computational burden; (2) iterative HMM searches using jackhmmer via the online webserver (89) in both directions (starting from the *Bombyx* or human/fly sequences) focusing on animal species, considering candidates that would be clearly part of single gene family; and (3) reciprocal HMM-vs-HMM profile searches with HHpred using *D. melanogaster* and *H. sapiens* and PFAM pre-calculated HMMs (31). If none of these methods provided any significantly similar candidates, they were not included in our analyses. For notes on gene discovery see Table S2.

Additional PsiBLAST (28), HMMER, and HHpred searches for BAT1 and WHAT were performed using the MPI Bioinformatics Toolkit (32, 92) (https://toolkit.tuebingen.mpg.de/) and the EMBL-EBI HMMER web server (93) (https://www.ebi.ac.uk/Tools/hmmer/home). Phylip Neighbor-joining tree of putative WHAT-like proteins was created with clustalw2 (94) without distance correction based on a multiple sequence alignment (MSA) generated with MUSCLE (95). MSA schematic was generated with Geneious Prime (Dotmatics).

## DATA AVAILABILITY STATEMENT

Stowers Institute original data underlying this manuscript can be accessed from the Stowers Original Data Repository at https://www.stowers.org/research/publications/LIBPB-2548. Sequencing data is submitted to the SRA and can be found under accession numbers SRR34018377-SRR34018405.

