## Supplemental figures and legends for "Programmed meiotic errors facilitate dichotomous sperm production in the silkworm, *Bombyx mori*"

### Supplementary Figure 1. Telomere clustering and SC formation in eupyrene spermatogenesis in *B. mori*.

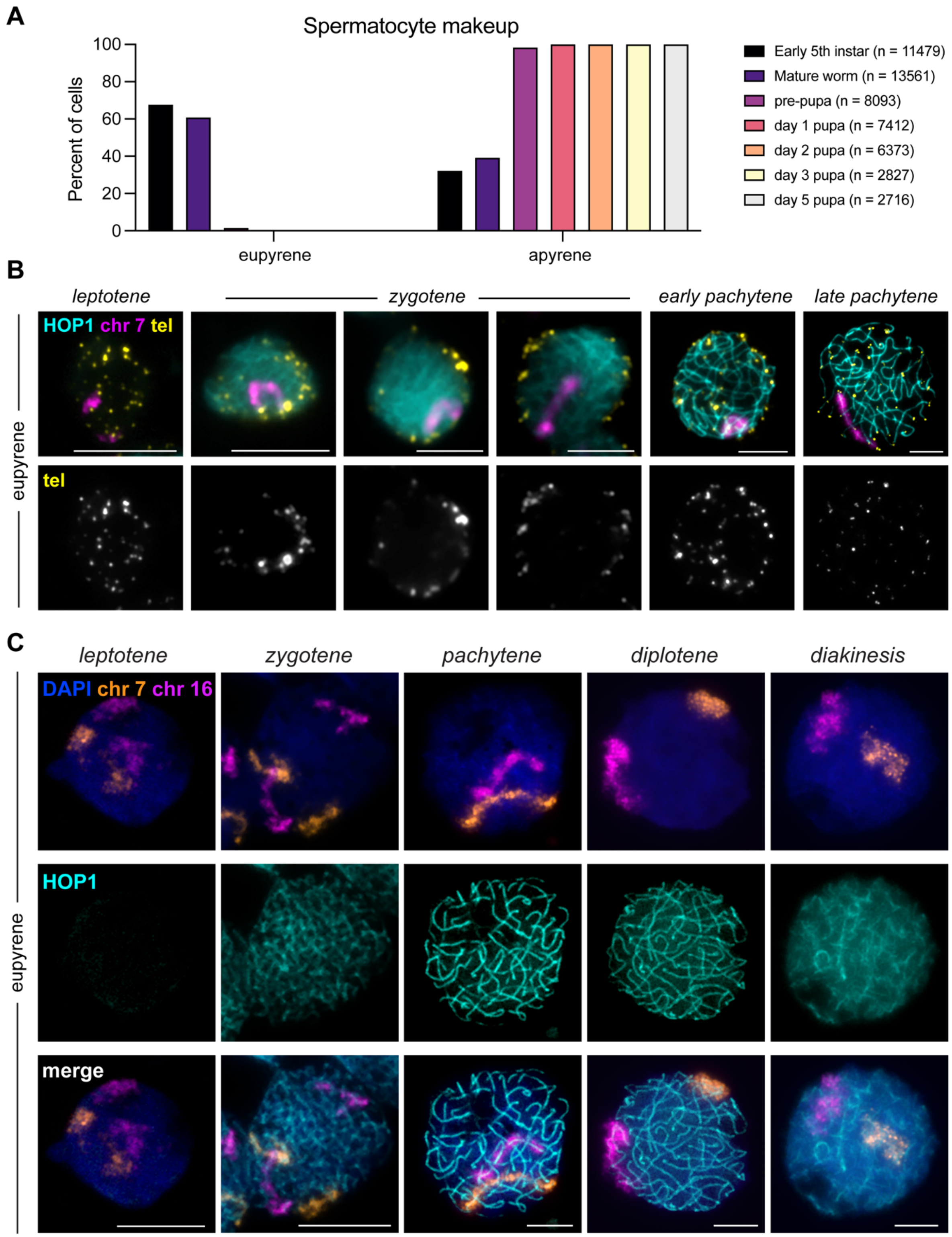

A. Quantification of eupyrene and apyrene spermatocytes throughout development. Using SYCP2 to distinguish between eupyrene and apyrene spermatocytes, the ratio between eupyrene and apyrene spermatocytes as a percentage of each cell type was calculated for the stages 5$^{th}$ instar, mature instar, pre-pupa, day 1 pupa, day 2 pupa, day 3 pupa, and day 5 pupa. In 5$^{th}$ and mature instars, the majority of cells are eupyrene spermatocytes (68% in 5$^{th}$ instar and 61% in mature instars). In pre-pupa, the majority are apyrene spermatocytes (98%). At remaining stages, no eupyrene spermatocytes were found. 5$^{th}$ instar, n = 11479. Mature instars, n = 13561. Pre-pupa, n = 8093. Day 1 pupa, n = 7412. Day 2 pupa, n = 6373. Day 3 pupa, n = 2827. Day 5 pupa, n = 2716. Counting was done using a mechanical cell counter. All counting was done on prophase I cells. Data came from cryosectioned slides.

B. Representative cells from 5$^{th}$ instar larval testes (eupyrene meiosis) spreads showing HOP1 (lateral element; cyan), Oligopaints for chr 7 (magenta), and telomere repeat FISH probes (yellow). As homologs align at zygotene, telomeres concurrently cluster around the nuclear periphery before dispersing again at the end of pachytene. Scale bar is 5 µm.

C. Cells from 5$^{th}$ instar larval testes (eupyrene meiosis) chromosome spreads. Cells are labeled with HOP1 (cyan), Oligopaints for chr 7 (orange), and Oligoppaints for chr 16 (magenta). DAPI is shown in blue. HOP1 localization to chromosomes begins in early zygotene (second column) and remains chromosome-associated throughout prophase I. Scale bar is 5 µm.

**Supplementary Figure 2. Apryene spermatogenesis in *B. mori* has aberrant SC formation.**

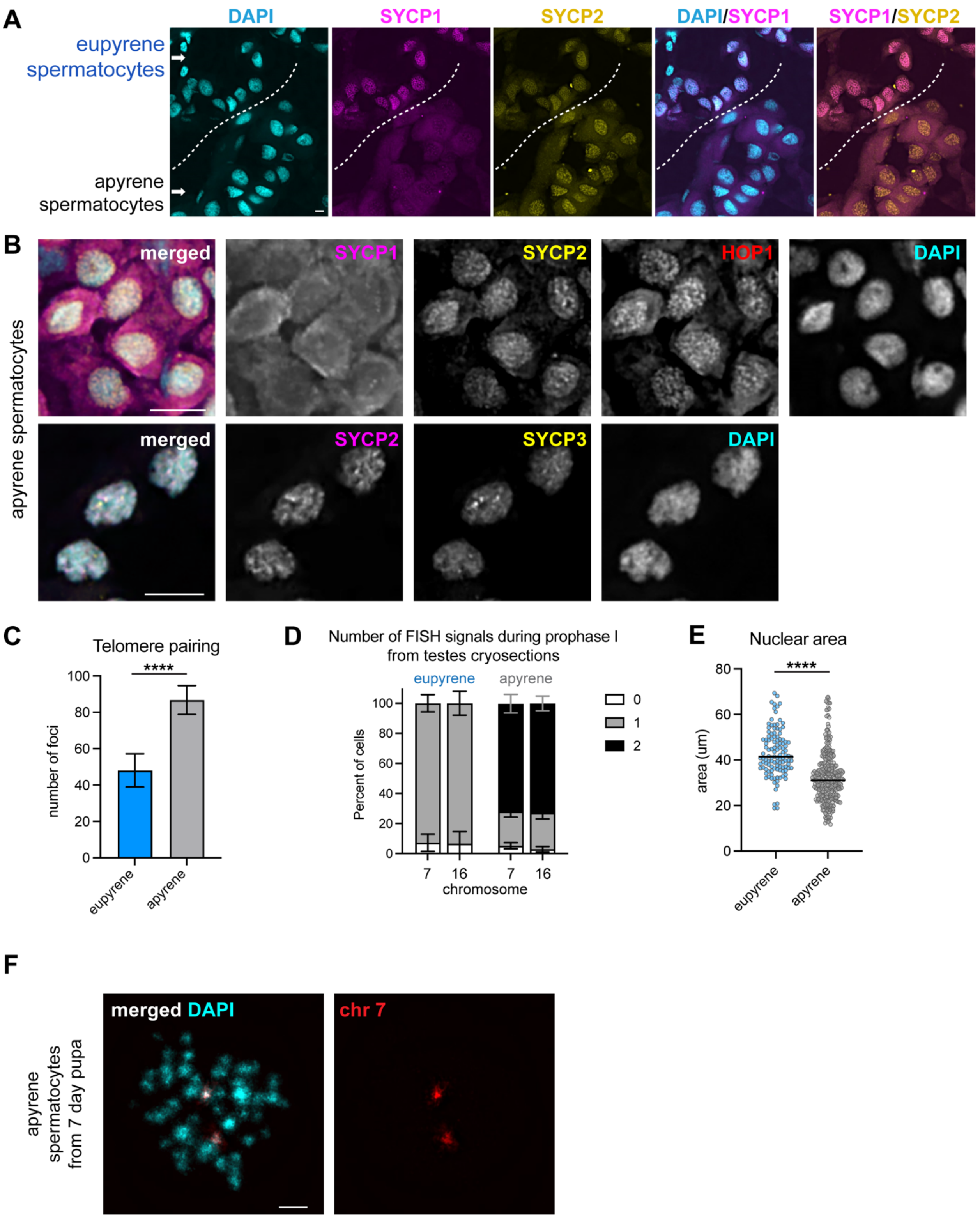

A. In mature larvae, both eupyrene and apyrene spermatocytes are detected. Apyrene spermatocytes show no SYCP1(magenta) and only short threads of SYCP2 (yellow). Eupyrene spermatocytes have full length threads of SYCP1(magenta) and SYCP2 (yellow). DAPI is shown in cyan. Scale bar, 5 μM. Images are maximum-intensity projections of the deconvolved *z*-series though the selected nuclei. *Associated with Figure 1.*

B. Representative images of apyrene spermatocytes from day 1 pupal testes. Top row: DAPI (cyan), SYCP1 (magenta), SYCP2 (yellow), and HOP1 (red). Bottom row: DAPI (cyan), SYCP2 (magenta), and SYCP3 (yellow). HOP1, SYCP2, and SYCP3 are localized to chromatin, but SYCP1 antibody shows only weak, non-specific signal. SYCP2, SYCP3, and HOP1 appear to form short threads. Scale bar is 5 μM. Images are maximum-intensity projections of the deconvolved *z*-series though the selected nuclei. *Associated with Figure 1.*

C. Bar graph showing number of telomeric FISH signals per nucleus in eupyrene ($5^{th}$ instar larvae) and apyrene (day 4 pupa) spermatocytes. Average number of telomeres was 48.1 ± 9.1 for eupyrene spermatocytes (n = 79) and 86.8 ± 7.9 for apyrene spermatocytes (n = 68). P<0.0001; were unpaired t-test (Kolmogorov-Smirnov test) between distributions.

D. Quantification of the number of whole chromosome Oligopaint signals in prophase I cells. Data from eupyrene spermatocytes from $5^{th}$ and mature instars averaged together had 1 signal in 92.7% & 93.3% (chr 7 & chr 16) of all samples (n = 199). This indicates that eupyrene spermatocytes are paired, with the error coming from small deviations due to sectioning and counting. The percentage that had zero signal was 7.3% and 6.67% (chr 7 and chr 16, n = 199). None had 2 signals. Conversely, apyrene spermatocytes from pre-pupa, day 1 pupa, day 2 pupa, and day 3 pupa averaged together 2 signals in 71.6% & 72.8% (chr 7 & chr 16) of all samples (chr 7, n = 779, chr 16, n = 996). This indicates that these chromosomes are not paired. The percentage that had 1 signal was 22.9% & 24.2% (chr 7, n = 779, chr 16, n = 996). The percentage that had zero signal was 5.21% & 2.94% (chr 7, n = 779, chr 16, n = 996). The lack of signal is likely due to sectioning and counting through different z-sections. Error bars are standard deviation. Data came from sectioned slides.

E. Dot plot showing area of eupyrene nuclei (early $5^{th}$ instar) and apyrene nuclei (day 1-3 pupae). The nuclei of eupyrene prophase I spermatocytes are significantly bigger than apyrene prophase I spermatocytes. DAPI was used as a proxy to measure nuclear size. Average size of eupyrene $5^{th}$ instar nuclei was 43.13 μm (n = 116), which was statistically bigger than the average area of nuclei in apyrene spermatocytes (33.3 μm, n = 320), For area calculations, see materials and methods. P<0.0001; were unpaired t-test (Kolmogorov-Smirnov test) between distributions. Data came from sectioned slides.

F.  Metaphase I spreads from day 7 pupal testes showing that apyrene spermatocytes at metaphase I do not have 28 bivalents as they would in eupyrene spermatocytes, DAPI (cyan) and whole chromosome Oligopaint for chr 7 (red). Scale bar is 5 μm. Images are maximum-intensity projections of the deconvolved *z*-series though the selected nuclei.

**Supplementary Figure 3. Metaphase I spindles in apyrene spermatocytes have distinct morphology compared to spindles in eupyrene spermatocytes.**

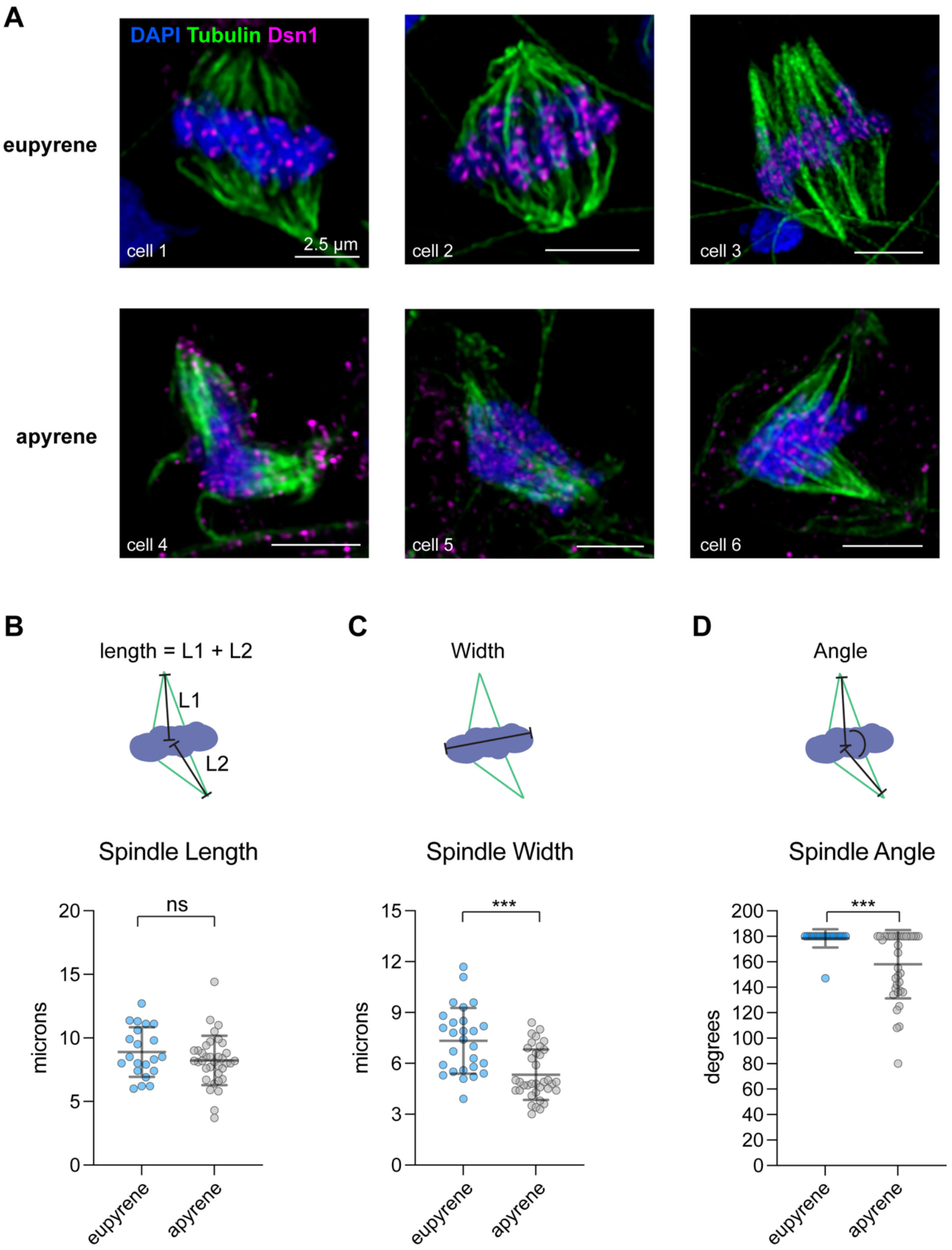

A. Representative images of IF on metaphase cells from eupyrene (top) and apyrene (bottom) testes chromosomes spreads. DAPI is shown in blue, tubulin in green, and Dsn1 in magenta. Scale bar is 2.5 µm. Spindles may appear slightly open due to loss of aster microtubules and centrosomes during the spreading process.

B - D. Top: Schematic of measurements taken for bottom graphs. Bottom: dot plot showing the spindle length (B), spindle width (C), and spindle angle (D) of eupyrene (blue) and apyrene (gray) metaphase I cells. ***$p \leq 0.0001$. Unpaired t-test with Welch's correction.

**Supplementary Figure 4. SYCP2, HOP1, and cohesin component REC8 remain on metaphase I chromosomes in eupyrene meiosis.**

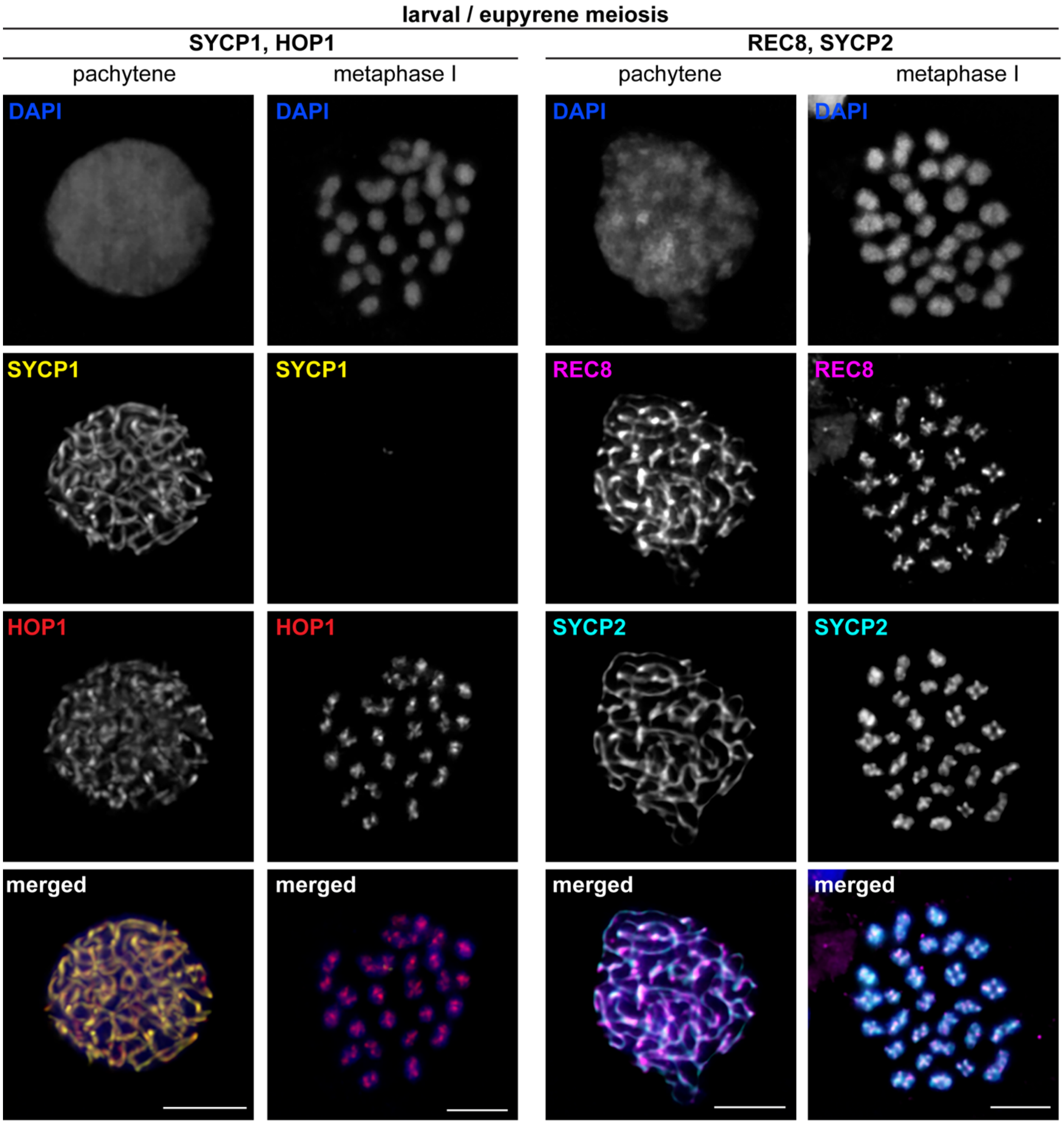


Representative images of IF on pachytene (left) and metaphase I (right) cells from larval testes chromosome spreads. Paired pachytene and metaphase I cells were in the same field of cells from the same slides. Left two columns: SYCP1 (yellow) and HOP1 (red). Right two columns: REC8 (magenta) and SYCP2 (cyan). HOP1, REC8, and SYCP2 remain on chromosomes through metaphase I in eupyrene meiosis. DAPI is shown in blue. Scale bar is 5 μm.

**Supplementary Figure 5. SYCP2, HOP1, and cohesin component REC8 remain on metaphase I chromosomes in apyrene meiosis.**

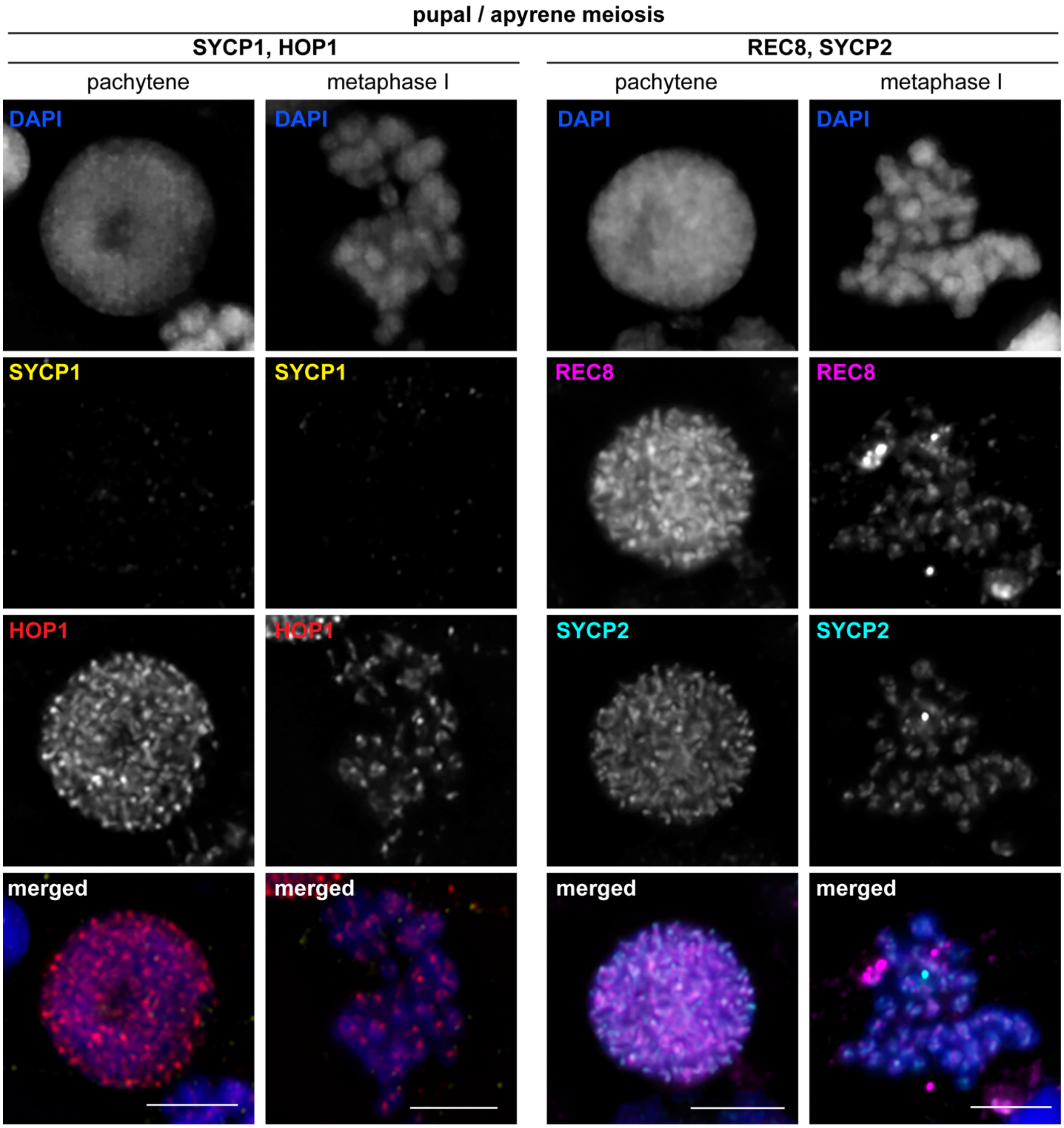


Representative images of IF on pachytene (left) and metaphase (right) cells from pupal testes chromosome spreads. Paired pachytene and metaphase cells were in the same field of cells from the same slides. Left two columns: SYCP1 (yellow) and HOP1 (red). Right two columns: REC8 (magenta) and SYCP2 (cyan). HOP1, REC8, and SYCP2 remain chromatin-associated through metaphase I in apyrene meiosis. DAPI is shown in blue. Scale bar is 5 μm.

**Supplementary Figure 6. Cell division gene expression comparison in larval testes and larval ovaries, and late-pupal testes.**

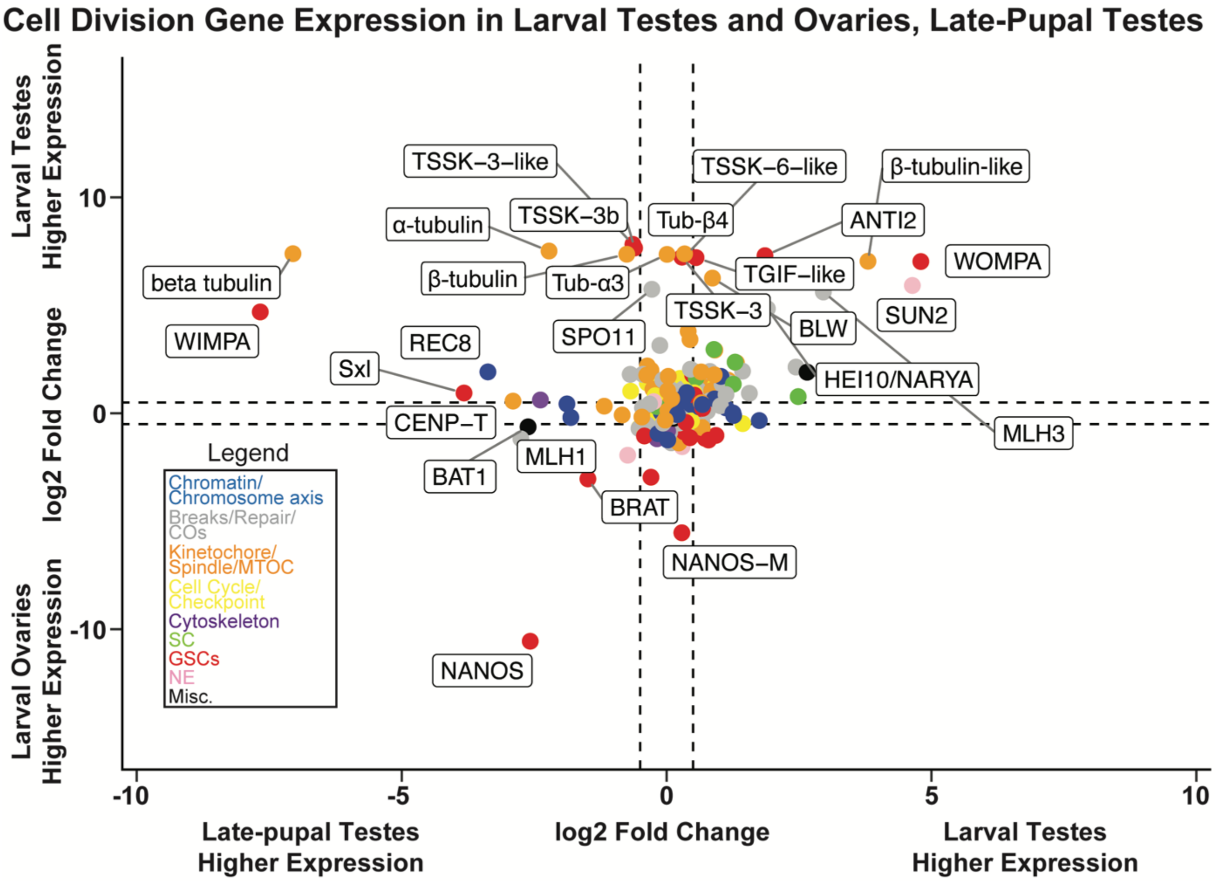


RNA-seq log2 fold change values of cell division genes between WT larval and late-pupal testes (x-axis) and WT larval testes and WT larval ovaries (y-axis). Select genes and their cell cycle roles, chromatin/chromosome axis (blue), breaks/repair/COs (gray), kinetochore/spindle/MTOC (orange), cell cycle/checkpoint (yellow), cytoskeleton (purple), synaptonemal complex (SC, green), GSCs (germline stem cells, red), nuclear envelope (NE, pink) and miscellaneous (black) are indicated in the legend.

**Supplementary Figure 7.** ***Sxl*** **gene expression tracks and MA plot.**

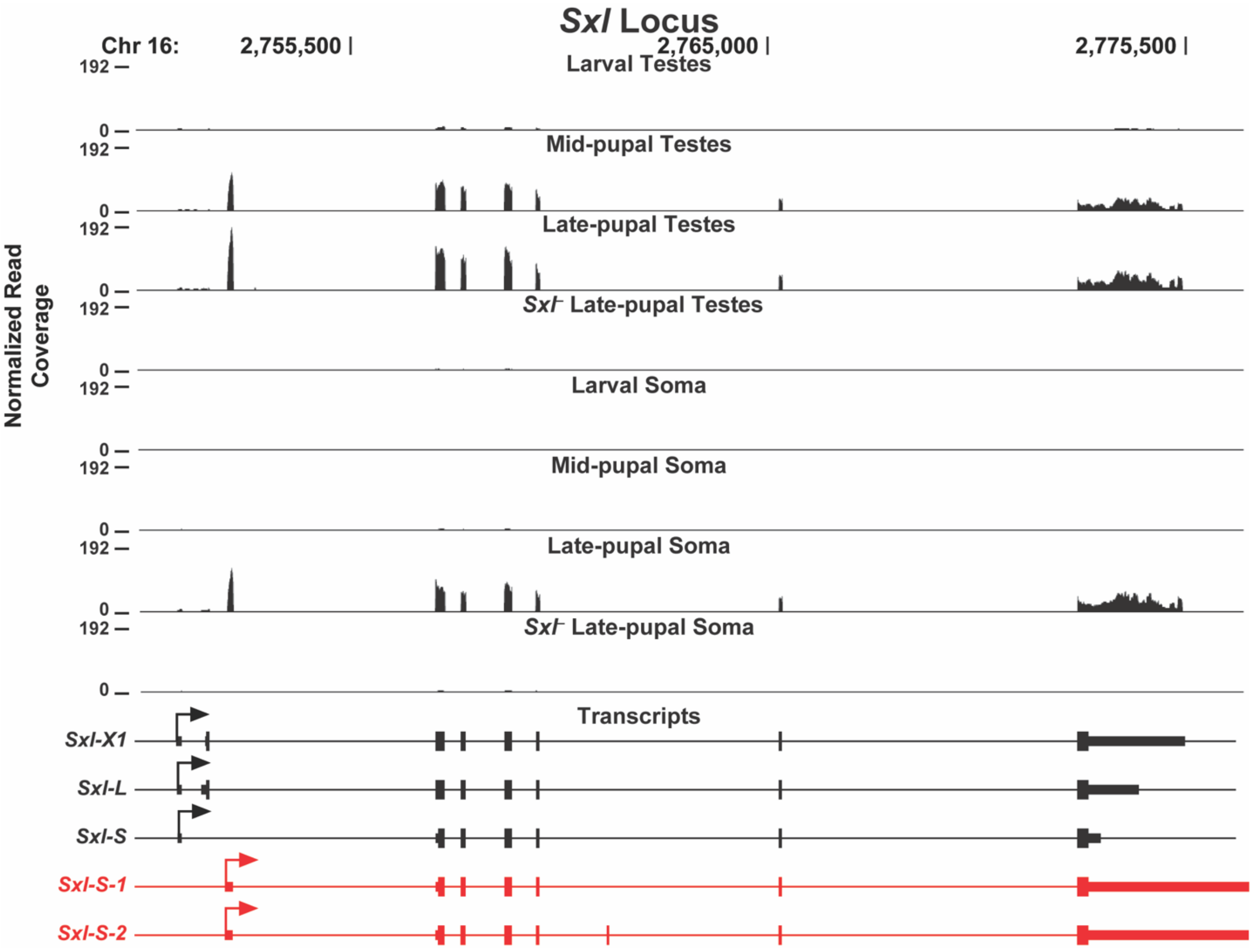


RNA-seq normalized read coverage tracks of the *B. mori sxl* locus for the indicated tissues and developmental timepoints. Arrows indicate transcriptional start sites, small rectangles indicate untranslated regions, large rectangles indicate open reading frames. Black transcripts represent NCBI annotated transcripts and red transcripts indicate *de novo* assembled transcripts from our analysis.

**Supplementary Figure 8. WHAT is a putative actin binding protein involved in *B. mori* eupyrene meiosis.**

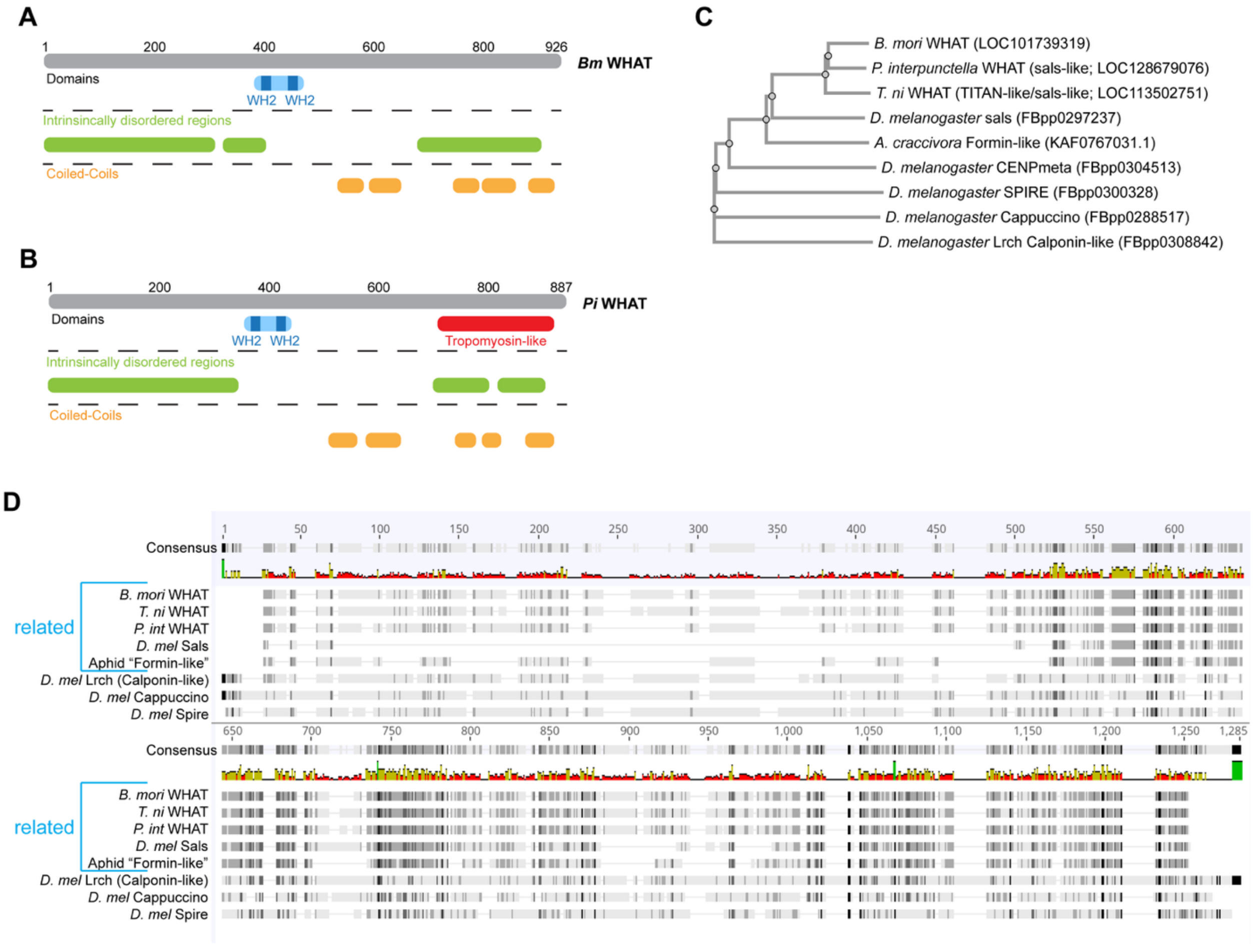


A. pfam protein domains identified in *B. mori* WHAT.

B. pfam protein domains identified in *P. interpunctella* WHAT.

C. Neighbor-joining tree of putative WHAT-like proteins in moths and flies, with one putative ortholog identified in aphids.

D. Multiple sequence alignment showing conservation, where darker color means more conserved. Based on the alignment, only the top 5 proteins are predicted to be orthologous.

**Supplementary Figure 9. Cell division gene expression comparison between larval ovaries and larval testes and late-pupal testes.**

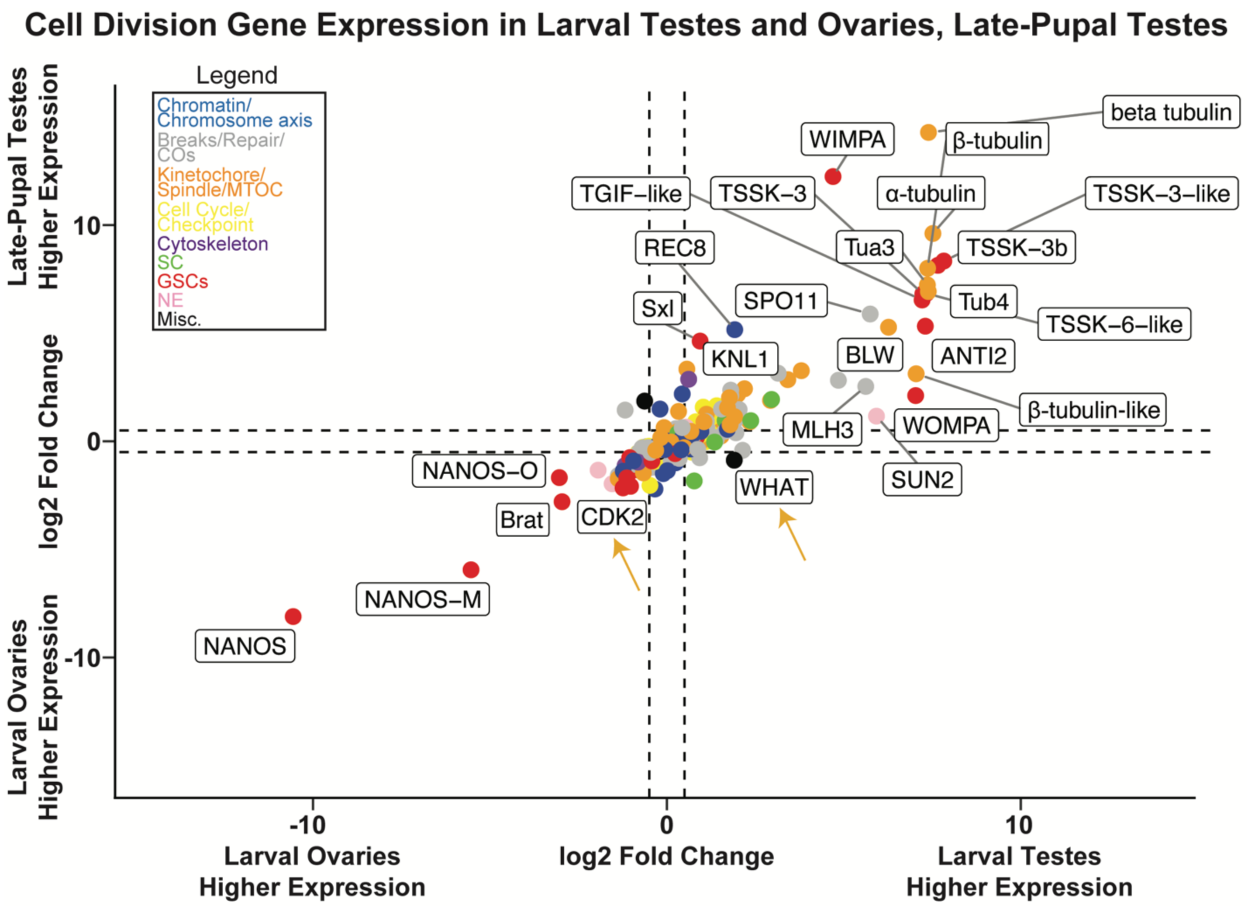


RNA-seq log2 fold change values of cell division genes between WT larval testes and larval ovaries (x-axis) and WT late-pupal testes and WT larval ovaries (y-axis). Select genes and their cell cycle roles, chromatin/chromosome axis (blue), breaks/repair/COs (gray), kinetochore/spindle/MTOC (orange), cell cycle/checkpoint (yellow), cytoskeleton (purple), synaptonemal complex (SC, green), GSCs (germline stem cells, red), nuclear envelope (NE, pink) and miscellaneous (black) are indicated in the legend. Arrows indicate genes CDK2 and WHAT as discussed in the text, which are more highly expressed in larval ovary than pupal testes

**Supplementary Figure 10. RNA-seq hierarchical replicate clustering and PCA plots.**

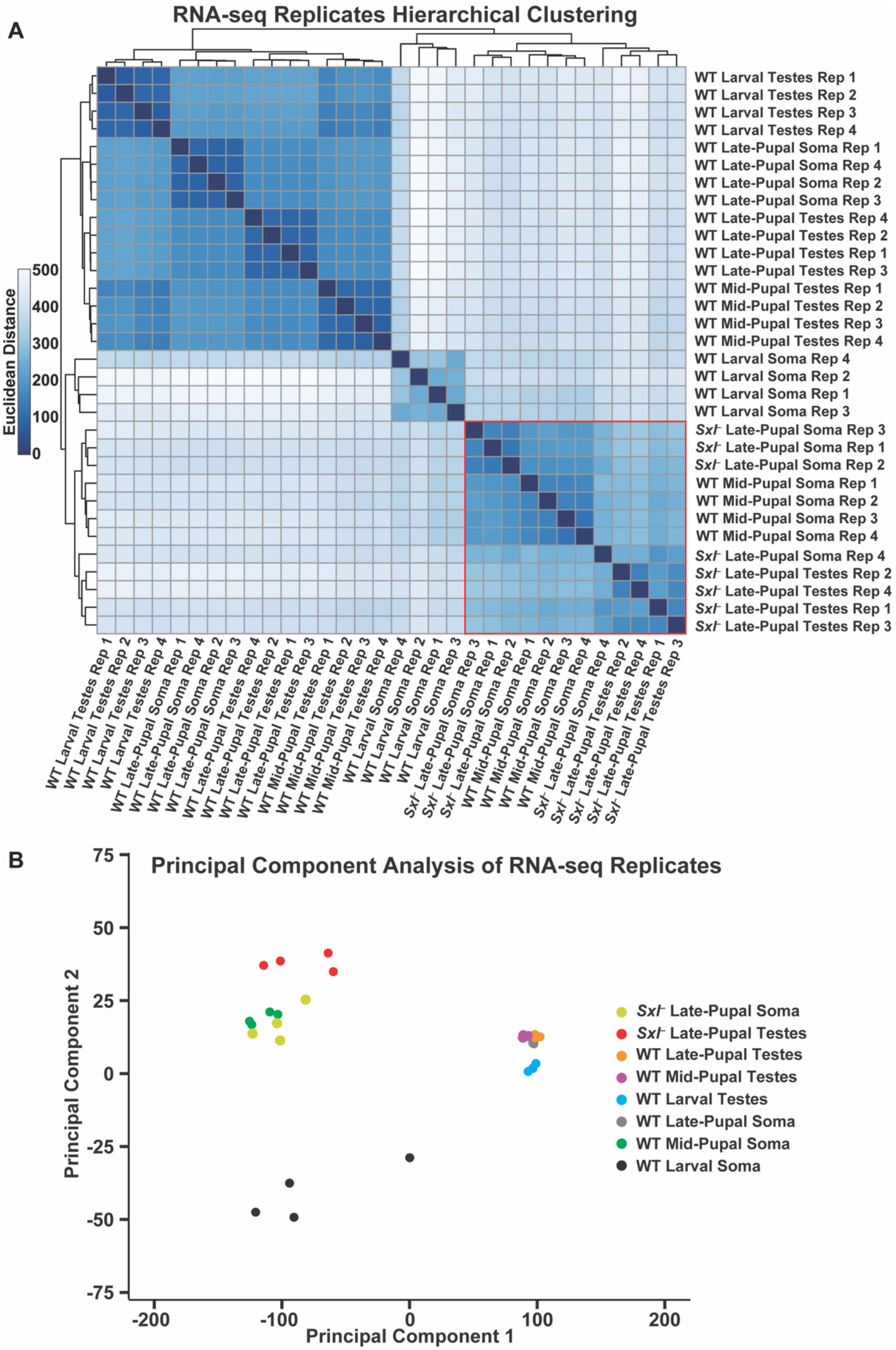

A. Heatmap measuring Euclidean distances of the respective RNA-seq replicates used in our analysis. Red box shows the location of the WT mid-pupal soma and *Sxl*- late-pupal testes samples.

B. Principal component analysis plot of *Sxl*- late-pupa soma (yellow), *Sxl*- late-pupa testes (red), WT late-pupal testes (orange), WT mid-pupal testes (purple), WT larval testes (blue), WT late-pupal soma (gray), WT mid-pupal soma (green), and WT larval soma (black) RNA-seq replicates used in our analysis.

**Supplementary Figure 11. Putative transcription factor expression between WT late-pupal testes and WT larval testes and *Sxl*- late-pupal testes.**

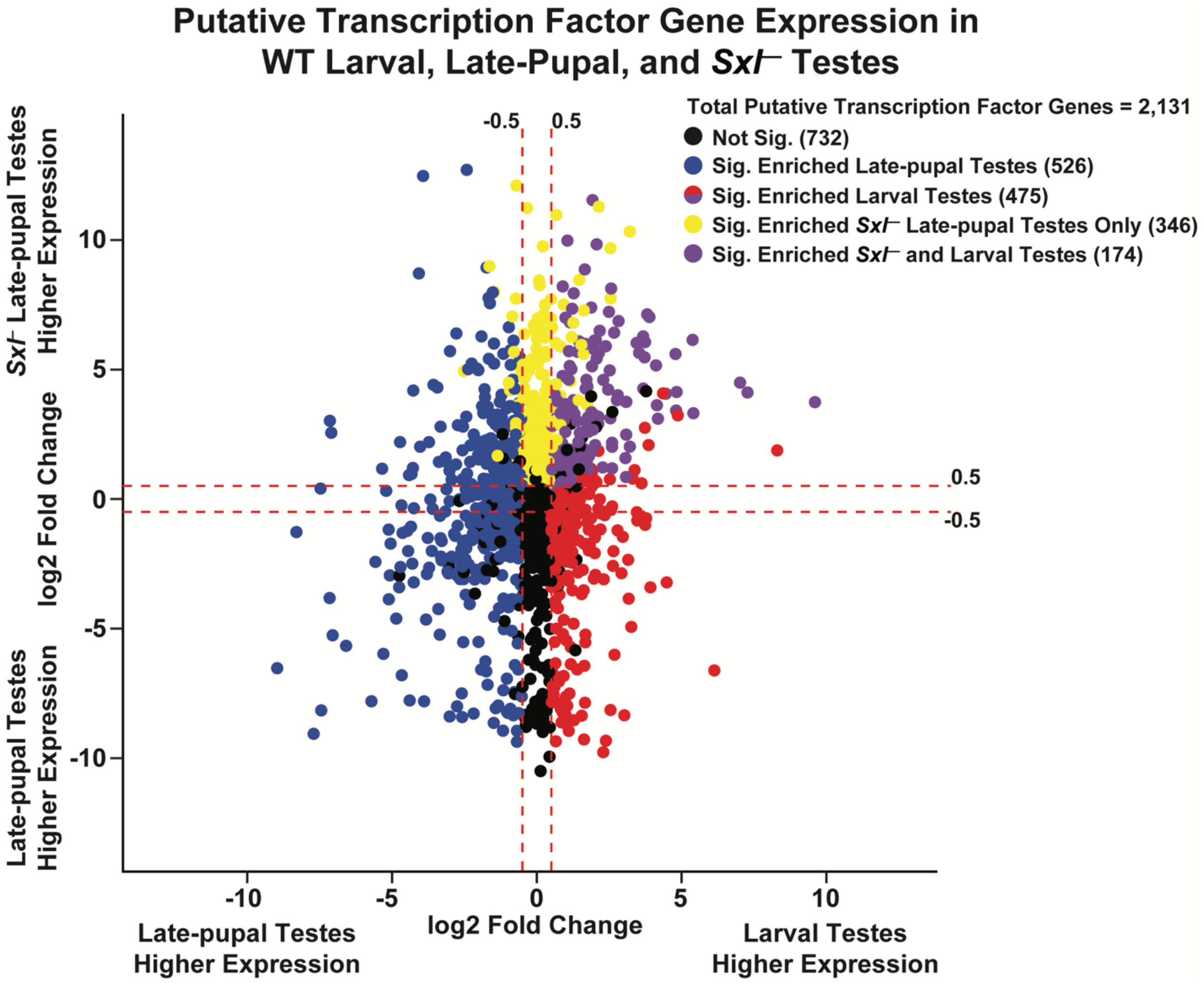


RNA-seq log2 fold change values of putative transcription factor genes between WT larval testes and WT late-pupal testes (x-axis) and *Sxl*- late-pupal testes and WT late-pupal testes (y-axis). Black dots indicate genes that are not differentially expressed. Blue dots indicate genes that are significantly higher expressed in WT late-pupal testes compared to WT larval testes. Red dots indicate genes that are significantly higher expressed in WT larval testes compared to WT late-pupal testes. Yellow dots indicate genes that are significantly higher expressed in *Sxl*- late-pupal testes compared to WT late-pupal testes. Purple dots indicate genes that are significantly higher expressed in both WT larval testes and *Sxl*- late-pupal testes compared to WT late-pupal testes.

**Supplementary Figure 12. Relative gene expression heatmaps of annotated hormone signaling genes.**

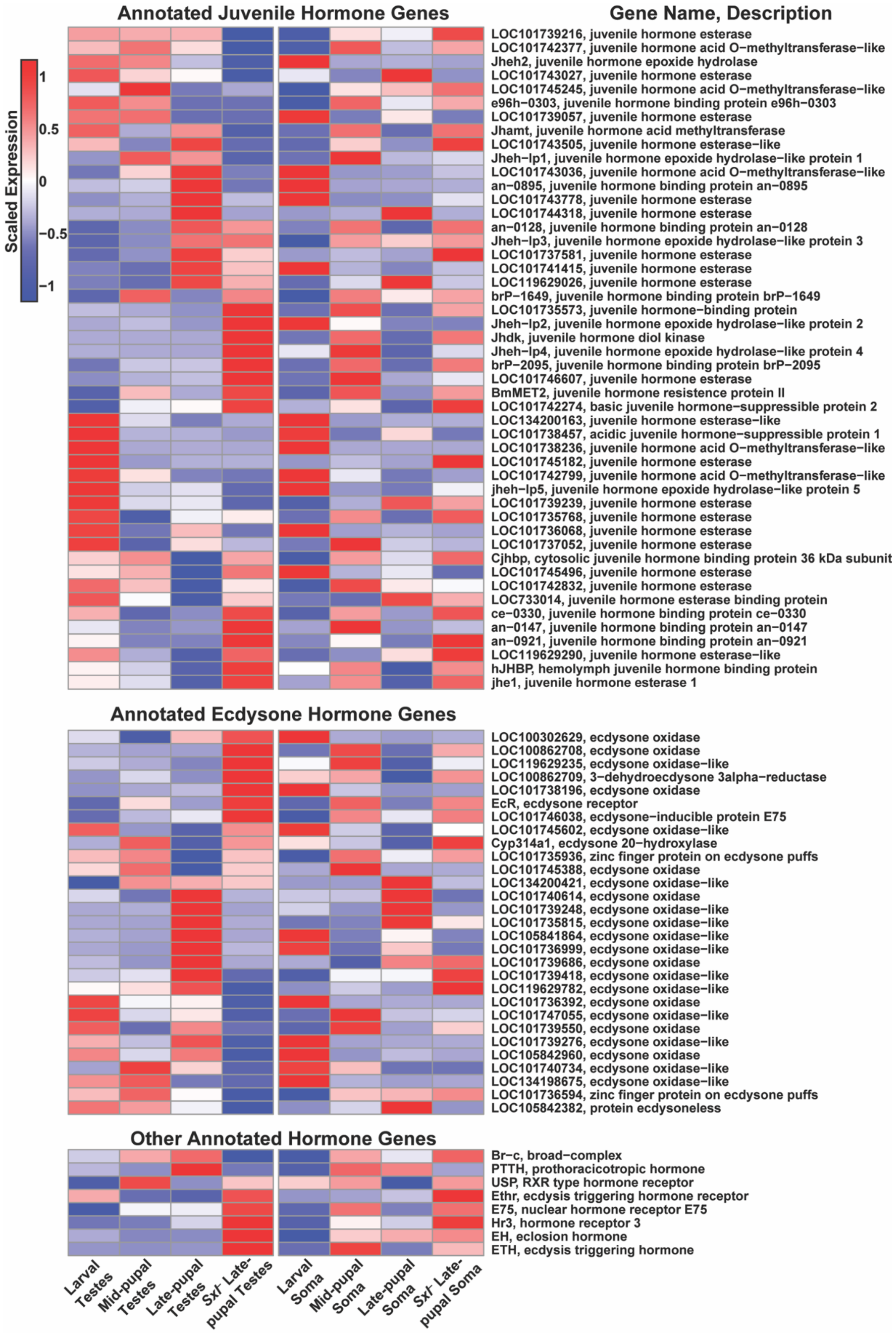
Annotated Juvenile Hormone Genes
Gene Name, Description
Scaled Expression
1
0.5
0
−0.5
−1
LOC101739216, juvenile hormone esterase
LOC101742377, juvenile hormone acid O−methyltransferase−like
Jheh2, juvenile hormone epoxide hydrolase
LOC101743027, juvenile hormone esterase
LOC101745245, juvenile hormone acid O−methyltransferase−like
e96h−0303, juvenile hormone binding protein e96h−0303
LOC101739057, juvenile hormone esterase
Jhamt, juvenile hormone acid methyltransferase
LOC101743505, juvenile hormone esterase−like
Jheh−lp1, juvenile hormone epoxide hydrolase−like protein 1
LOC101743036, juvenile hormone acid O−methyltransferase−like
an−0895, juvenile hormone binding protein an−0895
LOC101743778, juvenile hormone esterase
LOC101744318, juvenile hormone esterase
an−0128, juvenile hormone binding protein an−0128
Jheh−lp3, juvenile hormone epoxide hydrolase−like protein 3
LOC101737581, juvenile hormone esterase
LOC101741415, juvenile hormone esterase
LOC119629026, juvenile hormone esterase
brP−1649, juvenile hormone binding protein brP−1649
LOC101735573, juvenile hormone−binding protein
Jheh−lp2, juvenile hormone epoxide hydrolase−like protein 2
Jhdk, juvenile hormone diol kinase
Jheh−lp4, juvenile hormone epoxide hydrolase−like protein 4
brP−2095, juvenile hormone binding protein brP−2095
LOC101746607, juvenile hormone esterase
BmMET2, juvenile hormone resistence protein II
LOC101742274, basic juvenile hormone−suppressible protein 2
LOC134200163, juvenile hormone esterase−like
LOC101738457, acidic juvenile hormone−suppressible protein 1
LOC101738236, juvenile hormone acid O−methyltransferase−like
LOC101745182, juvenile hormone esterase
LOC101742799, juvenile hormone acid O−methyltransferase−like
jheh−lp5, juvenile hormone epoxide hydrolase−like protein 5
LOC101739239, juvenile hormone esterase
LOC101735768, juvenile hormone esterase
LOC101736068, juvenile hormone esterase
LOC101737052, juvenile hormone esterase
Cjhbp, cytosolic juvenile hormone binding protein 36 kDa subunit
LOC101745496, juvenile hormone esterase
LOC101742832, juvenile hormone esterase
LOC733014, juvenile hormone esterase binding protein
ce−0330, juvenile hormone binding protein ce−0330
an−0147, juvenile hormone binding protein an−0147
an−0921, juvenile hormone binding protein an−0921
LOC119629290, juvenile hormone esterase−like
hJHBP, hemolymph juvenile hormone binding protein
jhe1, juvenile hormone esterase 1
Annotated Ecdysone Hormone Genes
LOC100302629, ecdysone oxidase
LOC100862708, ecdysone oxidase
LOC119629235, ecdysone oxidase−like
LOC100862709, 3−dehydroecdysone 3alpha−reductase
LOC101738196, ecdysone oxidase
EcR, ecdysone receptor
LOC101746038, ecdysone−inducible protein E75
LOC101745602, ecdysone oxidase−like
Cyp314a1, ecdysone 20−hydroxylase
LOC101735936, zinc finger protein on ecdysone puffs
LOC101745388, ecdysone oxidase
LOC134200421, ecdysone oxidase−like
LOC101740614, ecdysone oxidase
LOC101739248, ecdysone oxidase−like
LOC101735815, ecdysone oxidase−like
LOC105841864, ecdysone oxidase−like
LOC101736999, ecdysone oxidase−like
LOC101739686, ecdysone oxidase
LOC101739418, ecdysone oxidase−like
LOC119629782, ecdysone oxidase−like
LOC101736392, ecdysone oxidase
LOC101747055, ecdysone oxidase−like
LOC101739550, ecdysone oxidase
LOC101739276, ecdysone oxidase−like
LOC105842960, ecdysone oxidase
LOC101740734, ecdysone oxidase−like
LOC134198675, ecdysone oxidase−like
LOC101736594, zinc finger protein on ecdysone puffs
LOC105842382, protein ecdysoneless
Other Annotated Hormone Genes
Br−c, broad−complex
PTTH, prothoracicotropic hormone
USP, RXR type hormone receptor
Ethr, ecdysis triggering hormone receptor
E75, nuclear hormone receptor E75
Hr3, hormone receptor 3
EH, eclosion hormone
ETH, ecdysis triggering hormone
Larval Testes
Mid-pupal Testes
Late-pupal Testes
Sxl- Late-pupal Testes
Larval Soma
Mid-pupal Soma
Late-pupal Soma
Sxl- Late-pupal Soma

Heatmap of annotated hormone gene expression (scaled RPKM values) in testes and somatic tissues across development. Genes are hierarchically clustered by their expression in testes tissues.

### Supplemental Table Legends

**Table S1. Meiosis gene expression in *B. mori* tissues.** log2 fold change values and p-adjusted values from differential expression analysis between *Sxl*- late-pupal vs WT late-pupal testes, WT larval vs WT late-pupal testes, and WT larval testes vs WT larval ovaries are listed for each meiotic gene.

**Table S2. All gene expression in *B. mori* tissues.** log2 fold change values and p-adjusted values from differential expression analysis between WT larval testes vs WT larval soma, WT late-pupal testes vs WT late-pupal soma, *Sxl*- late-pupal vs WT late-pupal testes, WT larval testes vs WT larval ovaries, and WT larval vs WT late-pupal testes are listed for each gene.

**Table S3. Significant GO terms for larval enriched gene expression.** Significant GO terms for significantly enriched gene expression in WT larval testes vs WT late-pupal testes.

**Table S4.** BAT1 psiBLAST and HHpred hits.

**Table S5.** WHAT psiBLAST and HHpred hits.

**Table S6. Gene level hierarchical cluster analysis.** Scaled mean RPKM values and cluster identity numbers for each gene in the cluster analysis.

**Table S7. Significant GO terms for cluster analysis.** Significant GO terms for genes belonging to each cluster from the hierarchical cluster analysis.

**Table S8. Putative transcription factors.** log2 fold change values and p-adjusted values from differential expression analysis between *Sxl*- late-pupal vs WT larval testes, WT larval vs WT late-pupal testes, and *Sxl*- late-pupal vs WT late-pupal testes are listed for each putative transcription factor.

**Table S9.** Antibodies and probes used in this study.
